# Sequence-independent substrate selection by the eukaryotic wobble base deaminase ADAT2/3 involves multiple protein domains and distortion of the tRNA anticodon loop

**DOI:** 10.1101/2022.06.16.496122

**Authors:** Luciano G. Dolce, Aubree A. Zimmer, Laura Tengo, Félix Weis, Mary Anne T. Rubio, Juan D. Alfonzo, Eva Kowalinski

## Abstract

The essential deamination of adenosine A_34_ to inosine at the wobble base is the individual tRNA modification with the greatest effects on mRNA decoding, empowering a single tRNA to translate three different codons. To date, many aspects of how eukaryotic deaminases specifically select their multiple substrates remain unclear. Here, using cryo-EM, we present the first structure of a eukaryotic ADAT2/3 deaminase bound to a full-length tRNA, revealing that the enzyme distorts the anticodon loop, but in contrast to the bacterial enzymes, selects its substrate via sequence-independent contacts of eukaryote-acquired flexible or intrinsically unfolded motifs distal from the conserved catalytic core. A novel gating mechanism for substrate entry to the active site is identified. Our multi-step tRNA recognition model yields insights into how RNA editing by A_34_ deamination evolved, shaped the genetic code, and directly impacts the eukaryotic proteome.

## Introduction

All nucleic acids in cells undergo post-transcriptional or post-replicative chemical modification, with tRNAs displaying the largest diversity of modified nucleotides. Modifications can affect the stability, folding and ultimately the function of tRNA in translation; in many cases RNA modification defects are associated with neurological disorders, cancers, or other diseases. In general, modifications in the core of a tRNA may sustain the canonical tRNA structure, while those occurring in the anticodon loop may fine-tune and streamline the essential process of protein synthesis at the ribosome (Björk et al., 1989; Grosjean and Westhof, 2016; Helm, 2006; Jonkhout et al., 2017; Motorin and Helm, 2022). Given the extraordinary degree of conservation in the 3D architecture of tRNAs, with their characteristic L-shape, it is fundamental to understand how different tRNA interacting proteins, including modification enzymes, can specifically recognize their cognate substrate tRNA and refuse structurally similar non-substrates. This is particularly critical for modifiers of the anticodon domain, as their action directly affects codon recognition and may influence the integrity of the cellular proteome (Bornelöv et al., 2019; Nedialkova and Leidel, 2015; Ranjan and Rodnina, 2016; Torres et al., 2021).

Several modifications occur at the first position of the anticodon (position 34) of tRNAs, the so-called wobble position, which base pairs to the third codon position of a base triplet in mRNAs (Grosjean and Westhof, 2016). Bacterial genomes do not contain any gene for a tRNA that per se can read the cytosine-ending (C-ending) codon triplet encoding arginine. Similarly, eukaryotes do not encode tRNAs that as such would read the C-ending codons for alanine, isoleucine, leucine, proline, serine, threonine, valine, and arginine (Chan and Lowe, 2016). To resolve this dilemma, a single tRNA (tRNA^Arg^_ACG_) in bacteria and 7-8 different tRNAs (depending on the organism) with an encoded A_34_ in eukaryotes (i.e. tRNA^Thr^_AGU_, tRNA^Ala^_AGC_, tRNA^Pro^_AGG_, tRNA^Ser^_AGA_, tRNA^Leu^_AAG_, tRNA^Ile^_AAU_, tRNA^Val^_AAC_, and tRNA^Arg^_ACG_) are post-transcriptionally modified to inosine (I_34_). This A_34_ to I_34_ tRNA deamination leads to the greatest enhancement in decoding capacity that can be caused by a single modification, as the INN anticodon can base pair with three different codon triplets at the mRNA, namely NNC (cytosine-ending), NNU (uracil-ending) and to a lesser extent NNA (adenosine-ending) (Agris et al., 2018; Crick, 1966; Dutta et al., 2022; Holley et al., 1965; Martin et al., 1985; Rafels-Ybern et al., 2019).

I_34_ formation is essential for viability and catalyzed by adenosine deaminases acting on tRNA (ADATs) (Gerber and Keller, 1999; Lyu et al., 2020; Rubio et al., 2007; Wolf et al., 2002). Several RNA-free bacterial tRNA deaminase A (TadA) structures were reported (Elias and Huang, 2005; Kim et al., 2006; Kuratani et al., 2005), followed by the *Staphylococcus aureus* enzyme bound to a synthetic hairpin mimicking the anticodon stem loop of bacterial tRNA^Arg^. TadA is a homodimeric enzyme, which binds one RNA ligand per protomer, forming a 2:2 protein : RNA complex. In this complex, the tRNA anticodon loop adopts an unusual conformation where the nucleosides are splayed out in a manner that is different from a canonical anticodon loop. TadA makes sequence-specific contacts to the exposed nucleobases (Losey et al., 2006). In eukaryotes, the essential enzyme catalyzing I34 formation in tRNA is a heterodimer, composed by one active (ADAT2) and one inactive (ADAT3) paralogous subunits, both homologs of the bacterial enzyme (Agris et al., 2018; Crick, 1966; Elias and Huang, 2005; Gerber and Keller, 2001; Sprinzl et al., 1998; Wolf et al., 2002). These ADATs carry a characteristic cytidine deaminase (CDA) active-site amino acid signature motif (C/H)XEX_n_PCXXC (with X being any amino acid, and n being any number of residues), containing the active site acidic glutamate (inert valine in inactive subunit ADAT3) and cysteine/histidine residues coordinating a Zn^2+^ cation (Bass, 2002; Betts et al., 1994; Kuratani et al., 2005). The motif, shared with other members of the CDA superfamily, is also found in the C-to-U deaminases which include the AID/APOBEC family.

In humans, mutations in the ADAT3 gene cause autosomal recessive rare disorders manifesting in a spectrum of intellectual disabilities and the tRNA pools isolated from affected individuals showed defects in A_34_-to-I_34_ deamination (Alazami et al., 2013; El-Hattab et al., 2016; Hengel et al., 2020; Ramos, 2020; Salehi Chaleshtori et al., 2018; Sharkia et al., 2019; Thomas et al., 2019). More recent investigations aimed to understand how the increased number of NNA codons in eukaryotic tRNAs pools and the expanded substrate recognition capacity by ADAT2/3 are related and how this could affect the proteome (Agris et al., 2018; Novoa et al., 2012; Torres et al., 2021). Furthermore, it has been reported that I_34_ in tRNA impacts gene expression regulation during differentiation of pluripotent stem cells by improving translation efficiency (Bornelöv et al., 2019).

Unlike bacterial TadA, which *in vitro* is also active on smaller harpin substrates, the eukaryotic ADAT2/3 requires a complete full-length tRNA L-shape for efficient deamination. Moreover, a classical 7-nucleotide anticodon loop has been shown to be a key determinant, since adding extra nucleotides to the loop interfered with I_34_ formation *in vitro* (Auxilien et al., 1996; Haumont et al., 1984). However, the structures of the ADAT2/3 enzymes in the absence of a tRNA ligand are not sufficient to explain how the tRNA architecture is recognized by ADAT2/3 (Liu et al., 2020; Ramos-Morales et al., 2021). Here we present the first structure of an ADAT deaminase bound to a full-length tRNA. Our cryo-electron microscopy (cryo-EM) structure of the tRNA-bound ADAT2/3 from *Trypanosoma brucei* (*T. brucei* or *Tb*) reveals a conserved distortion of the anticodon loop and exposure of the nucleotide bases in common with the bacterial catalytic pocket, with most contacts to the anticodon loop and the target nucleotide A_34_ provided by the active ADAT2 subunit. Through the evolution from a homo-to a heterodimeric configuration, both ADAT2/3 subunits have evolved additional peripheral binding regions, which provide sequence independent interactions that help to select and correctly position the tRNA substrate for catalysis. More generally, our findings provide insights into the evolution of these essential tRNA deamination enzymes.

## Results

### The cryo-EM structure and model of the *Trypanosoma brucei* ADAT2/3 - tRNA complex

To reveal how the eukaryotic tRNA deaminase (ADAT2/3) interacts with its substrate, we reconstituted the *T. brucei* protein-tRNA complex. Co-expression in insect cells and purification via a single 8-his-tag in *Tb*ADAT3 yielded a stoichiometric 1:1 *Tb*ADAT2-*Tb*ADAT3 complex and circumvented the enrichment of homodimers formed by the excess of *Tb*ADAT2 in the final sample, as *Tb*ADAT3 alone is insoluble (Extended Data Fig. 1a) (Rubio et al., 2007). Regardless of many attempts, the heterodimeric *Tb*ADAT2/3 complex failed to crystallize despite its extreme stability in stringent wash conditions (1 M KCl) during purification, and its elevated melting temperature of approximately 53.5°C in thermostability assays (Extended Data Fig. 1b). Co-purification of cellular tRNA^Thr^ and tRNA^Pro^ in initial purifications from *E. coli* (Extended Data Fig. 1c), led us to speculate that a tRNA indeed would stabilize the complex and create a particle size amenable for cryo-EM single-particle analysis. The 1:1:1 *Tb*ADAT2:*Tb*ADAT3:*Tb*tRNA^Thr^_CGU_ complex was reconstituted with *in vitro* transcribed *Tb*tRNA^Thr^_CGU_, a non-cognate tRNA omitting the reactive A_34_ that we chose to stabilize the complex by preventing product turnover (Extended Data Fig. 1d). The trimeric 95 kDa complex behaved well on quantifoil ultrAufoil gold grids and the preferred orientation of the particles was compensated by tilting the stage 30° for data collection. Data was processed with a combination of available software suites resulting in a final map with a resolution up to 3.6 Å, that is reasonable with respect to particle size (Extended Data Fig. 2 and Extended Data Table 1) (Punjani et al., 2017; Tegunov and Cramer, 2019). We rigid-body fitted the *Sc*tRNA^Phe^ (PDB: 1EHZ) but refined only the portion of the *Tb*tRNA^Thr^_CGU_ anticodon loop, which was well resolved (Extended Data Fig. 3a). The homologous yeast (PDB: 7BV5) and mouse (PDB: 7NZ7, 7NZ8) apo-crystal structures, and a model of the dimeric complex generated with AlphaFold served as templates for building and refinement of the heterodimeric *Tb*ADAT2/3 deaminase core (Extended Data Fig. 1e and 1f) (Jumper et al., 2021). The N-terminal domain of *Tb*ADAT3 (*Tb*ADAT3^N^) displayed poorer density, but a DeepEMhancer post-processed map allowed jiggle-fitting the AlphaFold model, without further refinement interventions (Sanchez-Garcia et al., 2021). The final model comprises the tRNA molecule, *Tb*ADAT2 residues 19-103, 123-175 and 183-221, and *Tb*ADAT3 residues 1-55, 105-263 and 277-340 (Fig. 1a and 1c).

**Fig. 1:**
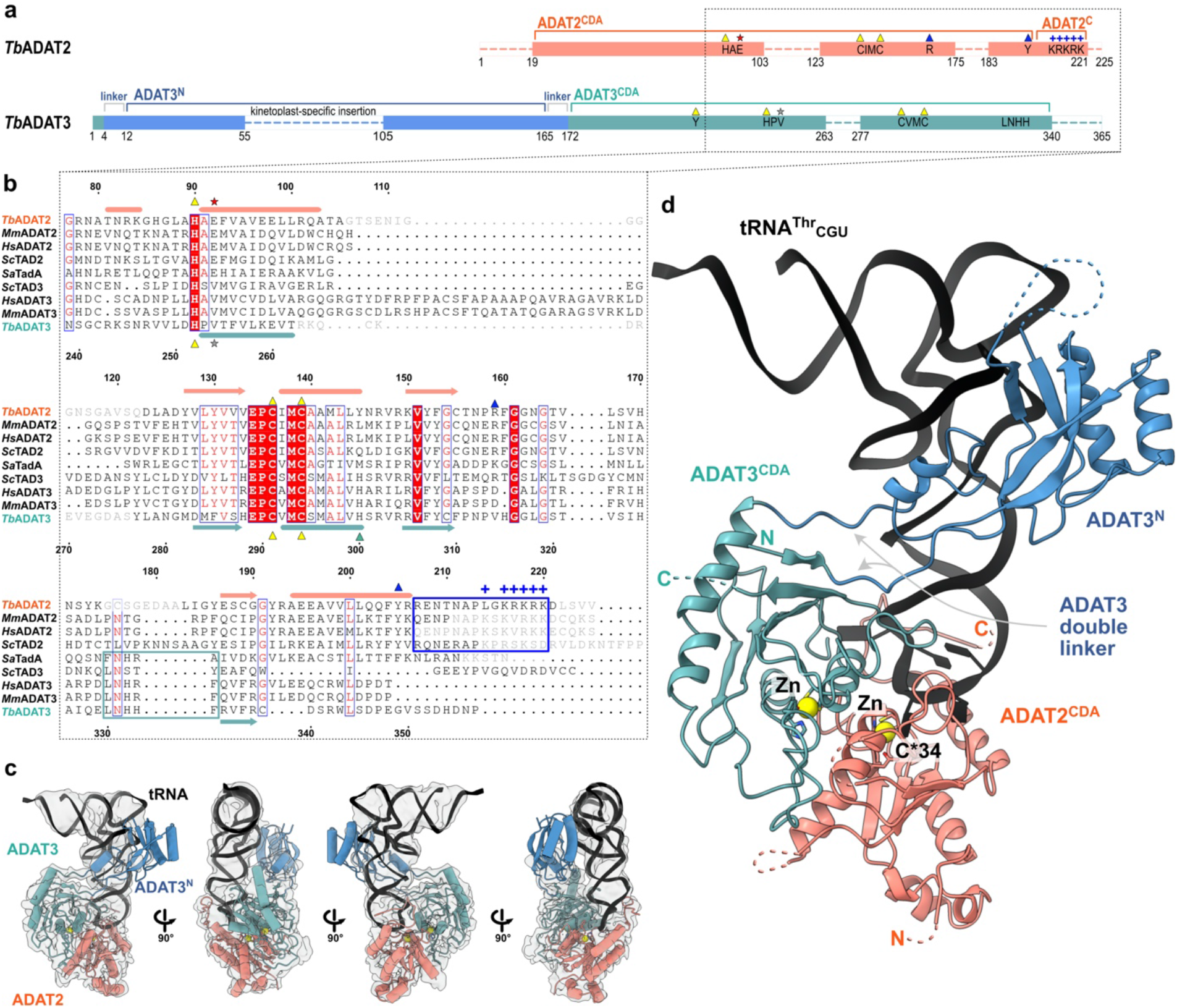
The structure of *Tb*ADAT2/3 heterodimer bound to tRNA^Thr^_CGU_. (**a**) Schematic representation of the protein subunits: ADAT2 CDA in salmon, ADAT3 N-terminal domain in blue, ADAT3^CDA^ in teal. Portions with traced atomic model as solid colors and non-resolved linkers as dotted lines. Important sequence motifs are annotated at their relative positions. Zn^2+^ coordinating residues are indicated by yellow triangles, the active site glutamate / pseudo active site valine by a red and gray star, ‘gate’ residues of ADAT2 with a blue triangle. Conserved positive charges of the KR-motif annotated as blue “+” signs. (**b**) Structural alignment of *Tb*, yeast and mouse ADAT2 and ADAT3 CDA domains and *Staphylococcus aureus* TadA. The sequences for human (*Hs*) ADAT2/3 included (*Mm*ADAT2/3 PDB: 7NZ7, *sc*TAD2/3 PDB:7 BV5, *sa*TadA PDB: 2B3J). Structural elements of *Tb* are annotated (ADAT2 above, ADAT3 below) with α-helices as cylinders and β-sheets as arrows. Zn^2+^ coordinating residues are indicated by yellow triangles, the active site glutamate / pseudo active site valine by a red and gray star, ‘gate’ residues of ADAT2 with a blue triangle and the active site histidine of ADAT3 with a teal triangle. ADAT2^C^ is boxed in blue with annotation of the conserved positive charges of the KR-motif (“+” signs). Grayed out letters indicate non-resolved loops in our structure and ADAT2^C^ residues not resolved in previous crystal structures. An ADAT3 conserved loop is annotated as a teal box. Other colors and boxes within the alignment represent relative conservation above 70%. (**c**) Four views of the final cryoSPARC map that was used for modeling (presented at a level of 0.09) with atomic model. Colors as above. (**d**) Cartoon model of ADAT2/3 bound to tRNA. Colors as in 1A. Zn^2+^ atoms in yellow, tRNA in black with only anticodon loop bases represented as slabs, the reactive C_34_ nucleotide is annotated as C*. Gray arrows point to the ADAT3 N-terminal domain double linker.

### The anticodon loop of the tRNA is remodeled upon ADAT2/3 binding

We inspected the overall conformation of the tRNA in the complex and noticed that the conformation of the anticodon-stem loop bound to the *Tb*ADAT2/3 core dramatically deviates from canonical free or ribosome A-site bound tRNA structures (Fig. 2a). In our complex structure, the *Tb*tRNA^Thr^_CGU_ is deeply inserted into the active site of *Tb*ADAT2 with all seven nucleotides (32-38) of the anticodon loop interacting directly with the ADAT2/3 deaminase core (Fig. 2b). While the stem nucleotides strictly obey Watson-Crick base pairing, and the non-canonical base transition pair between C_32_ and A_38_ is maintained (Auffinger and Westhof, 1999), the loop nucleotides 33-37 are extensively remodeled. Here, the sugar-phosphate backbone adopts a bent conformation and the nucleosides U_33_, C*_34_, G_35_ and A_37_ are splayed outwards (Fig. 2c and Extended Data Fig. 3a). The nucleobase of U_36_ is unconventionally internalized via a π-π stacking interaction between the nucleobase of C_32_ and residue Y205 of ADAT2 (Fig. 2d and Extended Data Fig. 3a). The unusual ADAT2/3-bound anticodon loop conformation with exposed bases, similar to the TadA bound hairpin, was unexpected, because the eukaryotic deaminase does not rely on sequence read-out (Auxilien et al., 1996; Losey et al., 2006; Ramos-Morales et al., 2021).

**Fig. 2:**
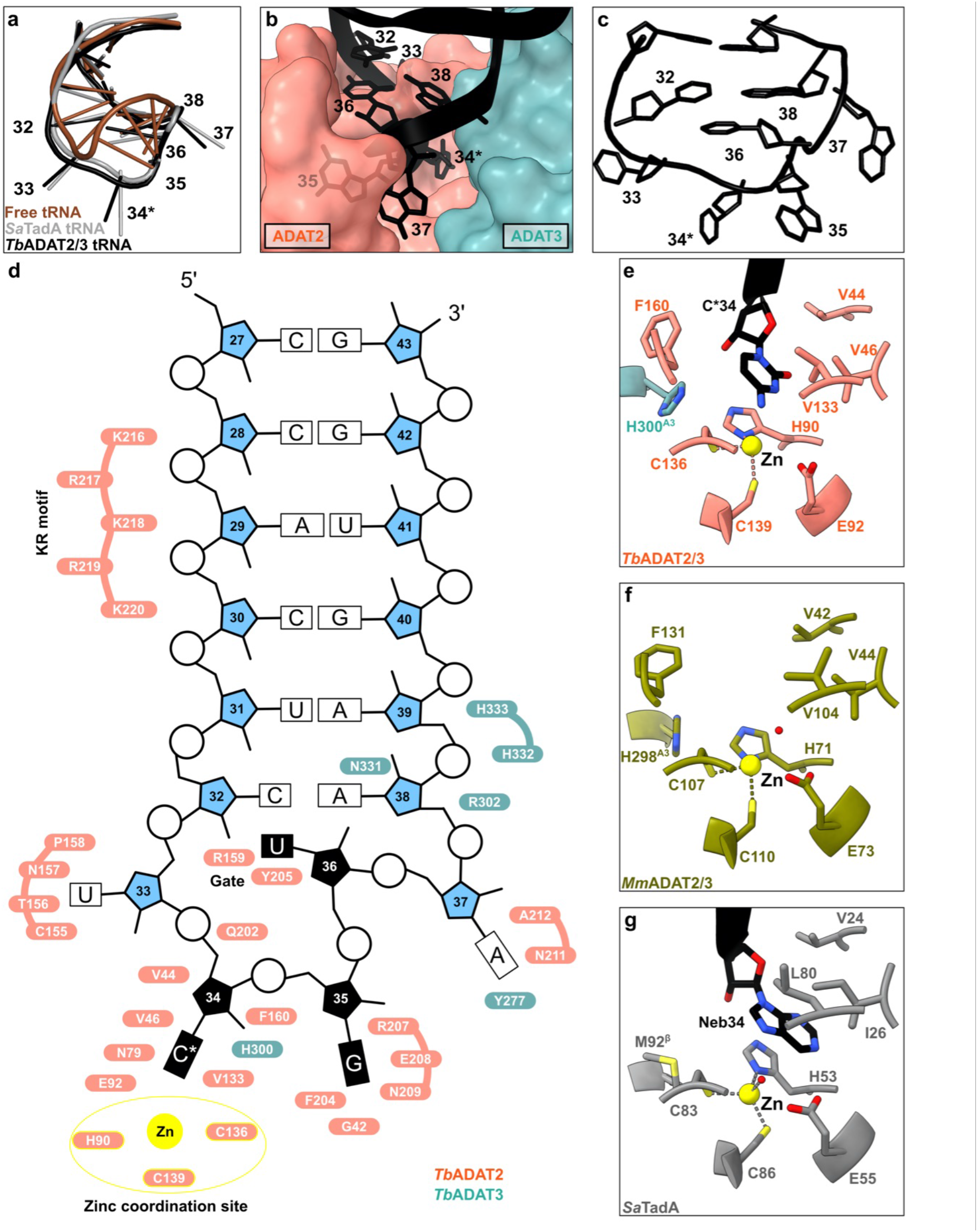
The interaction of the tRNA anticodon-stem loop with ADAT2/3. (**a**) The anticodon loop (ACL) bound to *Tb*ADAT2/3 (black) is distorted with respect to free tRNA (brown; PDB: 1EHZ) but resembles the ACL bound to bacterial TadA (grey; PDB: 2B3J), all cartoon-stick representation. (**b**) A deep cleft in the dimer interface accommodates the ACL. Colors as in figure 1, ADAT2/3 model as surface representation, tRNA as cartoon with anticodon loop bases shown. (**c**) The deformed ACL bound to the *Tb*ADAT2/3, backbone in cartoon and bases in stick representation. (**d**) Schematic overview of the ACSL and its interactions with *Tb*ADAT2/3. Zinc coordinating site highlighted in yellow. (**e, f, g**) Comparison of the active site pockets of *Tb* and *Mm*ADAT2/3 and TadA in the same orientation: (**e**) *Tb*ADAT2/3 with tRNA cytosine-base shown; (**f**) apo-*Mm*ADAT2/3 (PDB: 7NZ7), with the conserved H298^*Mm*ADAT3^ as the only ADAT3 residue present in the active site; (**g**) bacterial *Sa*TadA (PDB: 2B3J) with nebularine bound, with M92^*Sa*TadA**β**^ residue contributed by the second protomer of the homodimer. Residue side chains colored by heteroatom with zinc atoms in yellow.

### A deep catalytic pocket in ADAT2/3 accommodates A_34_

To determine the molecular basis for the unusual rearrangement of the tRNA, we examined how the binding pocket accommodates the anticodon loop. This pocket lies in the dimer interface between the cytidine deaminase (CDA) domains of *Tb*ADAT2 and *Tb*ADAT3. The flipped-out A_34_ (C_34_ in our construct) is deeply inserted into the active site of *Tb*ADAT2 (Fig. 2b). The active site of *Tb*ADAT2 features the typical CDA elements: a Zn^2+^ cation coordinated by two cysteines (C136^*Tb*ADAT2^ and C139^*Tb*ADAT2^) and a histidine (H90^*Tb*ADAT2^). Furthermore, the reactive nucleobase C*_34_ conformation is stabilized through hydrophobic interactions with V46^*Tb*ADAT2^ and V133^*Tb*ADAT2^ as well as hydrogen bonding with the conserved asparagine N79^*Tb*ADAT2^, while its ribose moiety is clamped by V44^*Tb*ADAT2^ and F160^*Tb*ADAT2^ (Fig. 2e and Extended Data Fig. 3b). The catalytic glutamic acid side chain, E92^*Tb*ADAT2^, and the catalytic water could not fully be resolved in our maps (Extended Data Fig. 3b). Besides these observations, the binding pocket for nucleotide A_34_ is remarkably similar between the tRNA-bound and -unbound structures, pointing to a certain rigidity of the active site to which the tRNA ligand must adapt (compare Figs. 2e, 2f, and Extended Data Fig. 3c). Remarkably, most residues in direct proximity to A_34_ are conserved between the bacterial and eukaryotic enzymes (Fig. 2g). An exception is the presence of histidine H300^*Tb*ADAT3^, which is conserved within ADAT3s (Fig. 1b, compare Figs. 2e and 2g). In summary, the central ADAT2/3 catalytic pocket for the C*_34_ nucleotide base is highly conserved between the bacterial homodimer and the eukaryotic heterodimer and no large conformational changes can be observed between the available eukaryotic substrate-bound and -unbound A_34_ pockets. Taken together these observations imply that for binding, the tRNA must adapt to ADAT2/3 through structural rearrangements.

### The anticodon loop is anchored by a molecular gate in ADAT2

In the bacterial enzyme TadA, extensive remodeling of the anticodon loop is provoked by favorable base-specific interactions, resulting in splayed-out anticodon loop nucleosides bound to a rather rigid protein surface (Losey et al., 2006). Since base-specific recognition does not take place in ADAT2/3 (Auxilien et al., 1996; Ramos-Morales et al., 2021), we looked for other features that could evoke such a non-canonical anticodon loop rearrangement. We identified a eukaryote-specific molecular “RY-gate” formed by residues arginine R159^*Tb*ADAT2^ and tyrosine Y205^*Tb*ADAT2^, which may provide a certain control mechanism for the tRNA to overcome before fully entering the catalytic pocket (Fig. 3). In the RNA-free *Mm*ADAT2/3 structures (PDB: 7NZ7, 7NZ8), this gate is closed through a cation-π interaction between these residues, R130^*Mm*ADAT2^ and Y171^*Mm*ADAT2^ (Fig 3a and Extended Data Fig. 3f). In the tRNA-bound *Tb*ADAT2/3 structure, the gate arginine R159^*Tb*ADAT2^ is released, and appears to be flexible as judged by the map density (Extended Data Fig. 3d). Opening of this gate also frees the side chain of the second gate residue Y205^*Tb*ADAT2^, which together with nucleobase C_32_, can sandwich and internalize nucleotide U_36_ via π-π stacking. This is likely to be a major driver in remodeling the anticodon loop (Extended Data Fig. 3e). Interestingly, despite the presence of an equivalent positively charged-aromatic pair (e.g. K106^*Sa*TadA^ and F145^*Sa*TadA^), this bacterial “gate” does not seem to operate because of suboptimal interacting geometry, which maintains these residues in an open conformation regardless of the presence or absence of RNA substrate; all available bacterial TadA structures in the PDB to date show an open configuration of these residues (Fig. 3b) (Dougherty, 2007; Gallivan and Dougherty, 1999; Kuratani et al., 2005; Losey et al., 2006). To test the importance of the gate residues, we measured *Tb*ADAT2/3 deamination activity of the wild type enzyme and several mutants. All amino acid substitutions that removed the aromatic nature of Y205^*Tb*ADAT2^ (namely Y205E and Y205R) had no detectable activity after 24-hour incubation, while a conservative mutation that maintains the aromaticity (Y205F) showed a reaction rate comparable to the wild-type enzyme. The necessity of the aromatic residue in this position indicates its importance in the base-sandwich for anticodon loop distortion. Next, we substituted the positive charge of the cationic gate residue R159^*Tb*ADAT2^ with a hydrophobic methionine (R159M), which led to an inactive enzyme. However, replacing R159^*Tb*ADAT2^ with tyrosine did not inactivate the enzyme but led to a 10-fold reduction of activity, indicating the importance of a positive charge for this gate residue. An “inverted gate” double mutant (R159Y, Y205R) was also inactive. A mutant mimicking the bacterial situation with a R159K substitution, strikingly, showed an almost 3-fold improved reaction rate compared to the wild type, suggesting a lysine in this position is more efficient for deamination activity (Figs. 3c, 3d, Extended Data Figs. 3j and 3k). Once it has entered the gate, the anticodon loop gains access to the deep joint anticodon binding pocket in order to establish further interactions with the ADAT2 core: asparagine N157^*Tb*ADAT2^ can form a hydrogen bond with the hydroxyl of the U_33_ ribose and the splayed-out nucleotide U_33_ can interact with ADAT2 residues 155^*Tb*ADAT2^-158^*Tb*ADAT2^ as well as glutamine Q202^*Tb*ADAT2^ (Fig. 2d). An ADAT3 RNA binding loop, 328-333^*Tb*ADAT3^, conserved between TadA and ADAT3 but not ADAT2, establishes contacts with the tRNA backbone at nucleotides 38-40 and the nucleobase A_38_ via two histidines (H332^*Tb*ADAT3^ and H333^*Tb*ADAT3^) and an asparagine N331^*Tb*ADAT3^, respectively (framed teal in Fig. 1b, Fig. 2d, Extended Data Figs. 3g, 3h, 3i). In conclusion, we identify a molecular gate formed by a cation-π interaction in eukaryotic ADAT2 which needs to be penetrated by the tRNA; the opened gate allows stabilizing aromatic stacking interactions with the C_32_ and U_36_ nucleobases to facilitate access to the deep anticodon loop binding cleft where further favorable protein-RNA interactions are established, altogether promoting the unusual anticodon loop configuration.

**Fig. 3:**
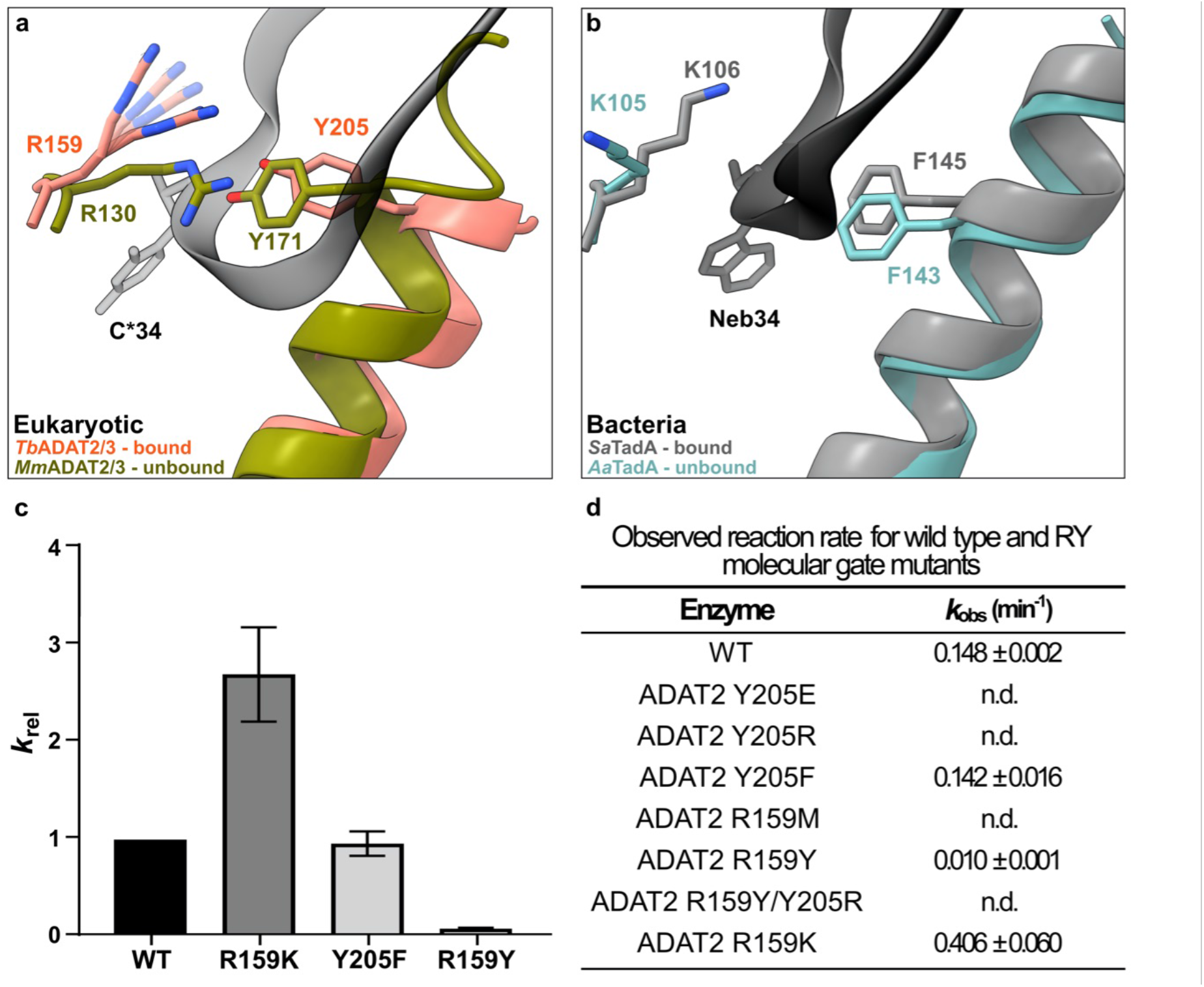
The tRNA anticodon loop passes through a molecular RY-gate in ADAT2. (**a**) Superposition of the tRNA-bound gate in *Tb*ADAT2/3 and the apo-gate in *Mm*ADAT2/3 (PDB: 7NZ7). The side chain of R159^*Tb*ADAT2^ is represented as flexible due to its poor side chain density. (**b**) The corresponding positions in bacteria deaminase TadA, RNA bound versus unbound (PDB: 2B3J and 1WWR respectively). (**c**) Fold change of the observed rate constants of mutants to wild-type (*k*_rel_ = *k*_obs_ mutant/*k*_obs_ WT). Graph values calculated from three independent replicates. (**d**) Observed reaction rate for wild type and RY molecular gate mutants. All mutants are of *Tb*ADAT2 co-expressed with *Tb*ADAT3. n.d. denotes no detectable activity after 24-hour assay incubation. *k*_obs_ values obtained from single-turnover kinetic assays (n=3)

### An intrinsically disordered motif in the ADAT2 C-terminus embraces the anticodon stem loop

The available crystal structures of apo-ADAT2/3 failed to resolve the very C-terminal portion of ADAT2 (ADAT2^C^, residues 207-225^*Tb*ADAT2^), a segment which is absent in bacterial TadA proteins. This region carries a conserved, positively charged motif at the C-terminal end of ADAT2^C^, herein referred to as “KR-motif” (residues 216-220^*Tb*ADAT2^ KRKRK) (Fig. 1b). The KR-motif was previously identified as essential for tRNA binding and deamination (Ragone et al., 2011). We fitted ADAT2^C^ into the cryo-EM map guided by the AlphaFold model of the complex; following the C-terminal helix which is the last element resolved in the crystal structures, a conserved proline induces a turn of the polypeptide, such that the KR-motif bends back towards the helix (Fig. 4). ADAT2^C^ displays extensive contacts with the anticodon loop and stem: residues 207-209^*Tb*ADAT2^ and 210-213^*Tb*ADAT2^ of *Tb*ADAT2^C^ embrace the flipped-out nucleobases G_35_ and A_37_ (Figs. 2d, 4a). Furthermore, the positively charged C-terminal KR-motif, which aligns into the major groove of the anticodon loop, likely provides extended interactions to the phosphate backbone of the anticodon stem towards nucleotides 28-31 (Figs. 4b, 4b). The less clear density here might reflect a certain fluidity of the interaction and adaptation to different sequence registers could be imagined, which would be required to adjust to the different types of tRNA. Taken together, ADAT2^C^ including the essential KR-motif seems to be intrinsically flexible in the absence of tRNA, but can reorganize and adjust to bind into the major groove of the anticodon loop and also establish interactions with portions of the anticodon stem.

**Fig. 4:**
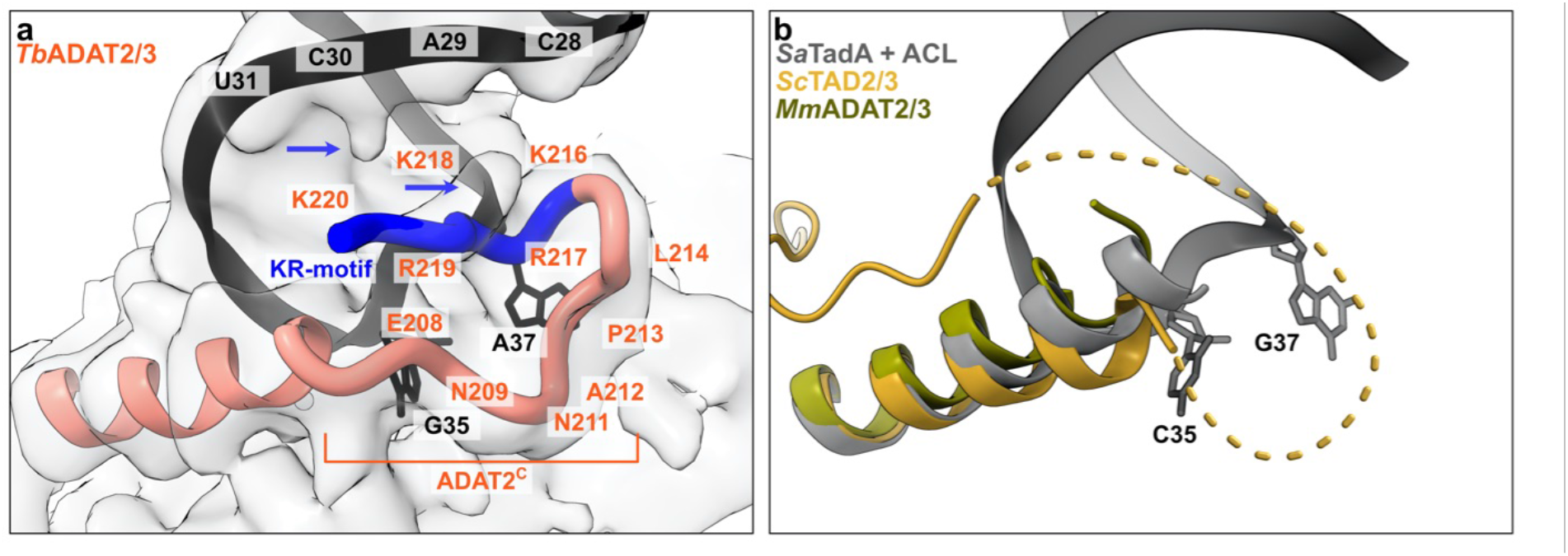
The ADAT2 C-terminal region including the KR-motif. (**a**) The C-terminus of ADAT2 embraces nucleotides 35 and 37 and KR-loop residues (K or R) provide additional contacts with the anticodon stem loop via phosphate backbone interactions at residues 28-31. A cartoon representation of the ADAT2 C-terminus and tRNA is shown with the cryo-EM map. Due to the limited resolution, we refrain from showing side chains in the ADAT2^C^ model but instead indicate map density representing protein-RNA contacts by blue arrows. (**b**) The same region of *Sc*TAD2/3 (PDB: 7BV5), *Mm*ADAT2/3 (PDB: 7NZ7) and RNA bound *Sa*TadA (PDB: 2B3J). The flexible unmodeled loop of *Sc*ADAT2 is shown as a dotted line.

### The N-terminal domain of *Tb*ADAT3 binds the tRNA via multiple contacts along the anticodon arm and elbow region

Experimental evidence has suggested a role of the eukaryotic ADAT3 N-terminal domain in RNA binding, which we seek out to elucidate in detail through the cryo-EM structure. AlphaFold and RoseTTA models (Anishchenko et al., 2021; Jumper et al., 2021) of ADAT3^N^ reveal a domain containing a slightly twisted 4-stranded antiparallel β-sheet with two α-helices in a βαββαβ arrangement (residues 13-140^*Tb*ADAT3^) and an additional appended α-helix (residues 141-164^*Tb*ADAT3^). This constellation closely resembles the crystal structures of the mouse and yeast homologues, with the main difference being a kinetoplast-specific insertion (residues 57-99^*Tb*ADAT3^) that is predicted to be disordered (Fig. 5a, Extended Data Fig. 1e and 4a). Snapshots collected from previous crystal structures placed this domain in various positions with respect to the deaminase domains, suggesting that this domain is flexibly attached to the catalytic core. However, all previously observed positions disagree with our cryo-EM map or would clash with other parts of the model. The ADAT3 N-terminal domain is connected to the protein core between two linkers (N-terminal: residues 4*-*12^*Tb*ADAT3^; C-terminal: residues 165-172^*Tb*ADAT3^) which form together a quasi-parallel double linker and are well-resolved in our maps (Fig. 1d). We jiggle-fitted the ADAT3^N^ AlphaFold model into the post-processed DeepEMhancer map without further side chain refinement (Extended Data Figs. 4b and 4c). In the model we identify two main positively charged contact regions between the ADAT3^N^ and the tRNA (Fig 5a). The first interaction region (spot ‘A’ in Fig. 5a) represents an elongated positively charged stretch facing the minor groove of the D-stem of the tRNA, likely binding the phosphate backbone via charged interactions. This region comprises the appendix α-helix of ADAT3^N^ together with the C-terminal linker that connects to the deaminase domain. A second interaction hotspot can be observed in the interface between the ADAT3^N^ domain and portions of the D-loop and elbow of the tRNA, in particular in the region of nucleotides 19 and 56, a base pair that is fundamental to the 3D structure of any tRNA (spot ‘B’ in Fig. 5a) (Levinger et al., 1998; Quigley and Rich, 1976). ADAT3 residues in both these regions have been attributed a role in tRNA binding and deaminase activity in previous experiments (Fig. 5b, Extended Data Fig. 4a and Extended Data Table 2) (Liu et al., 2020; Ramos-Morales et al., 2021). The earlier observation that long variable arms in tRNA^Val^ or tRNA^Ser^ do not affect inosine formation, is in accordance with our data as the map does not show density in the region where the variable arm would be positioned (Achsel and Gross, 1993; Auxilien et al., 1996; Ragone et al., 2011; Spears et al., 2011a). We reasoned that the elongated, continuous positive patch is putatively able to bind RNA via phosphate backbone interactions independently of sequence context. The somewhat lower resolution of ADAT3^N^ with respect to the deaminase core may indicate several possible binding modes and potential for adaptation to different tRNA molecules. The fact that ADAT2/3 is inactive on truncated anticodon loop substrates points towards the requirement of these extended interactions with ADAT3^N^ for correct insertion of the tRNA into the active site (Auxilien et al., 1996; Elias and Huang, 2005). In conclusion, our structure reveals a critical contribution of the ADAT3 N-terminal domain in the recognition of intact tRNA architectures via electrostatic contacts along the anticodon arm and in the elbow region.

**Fig. 5:**
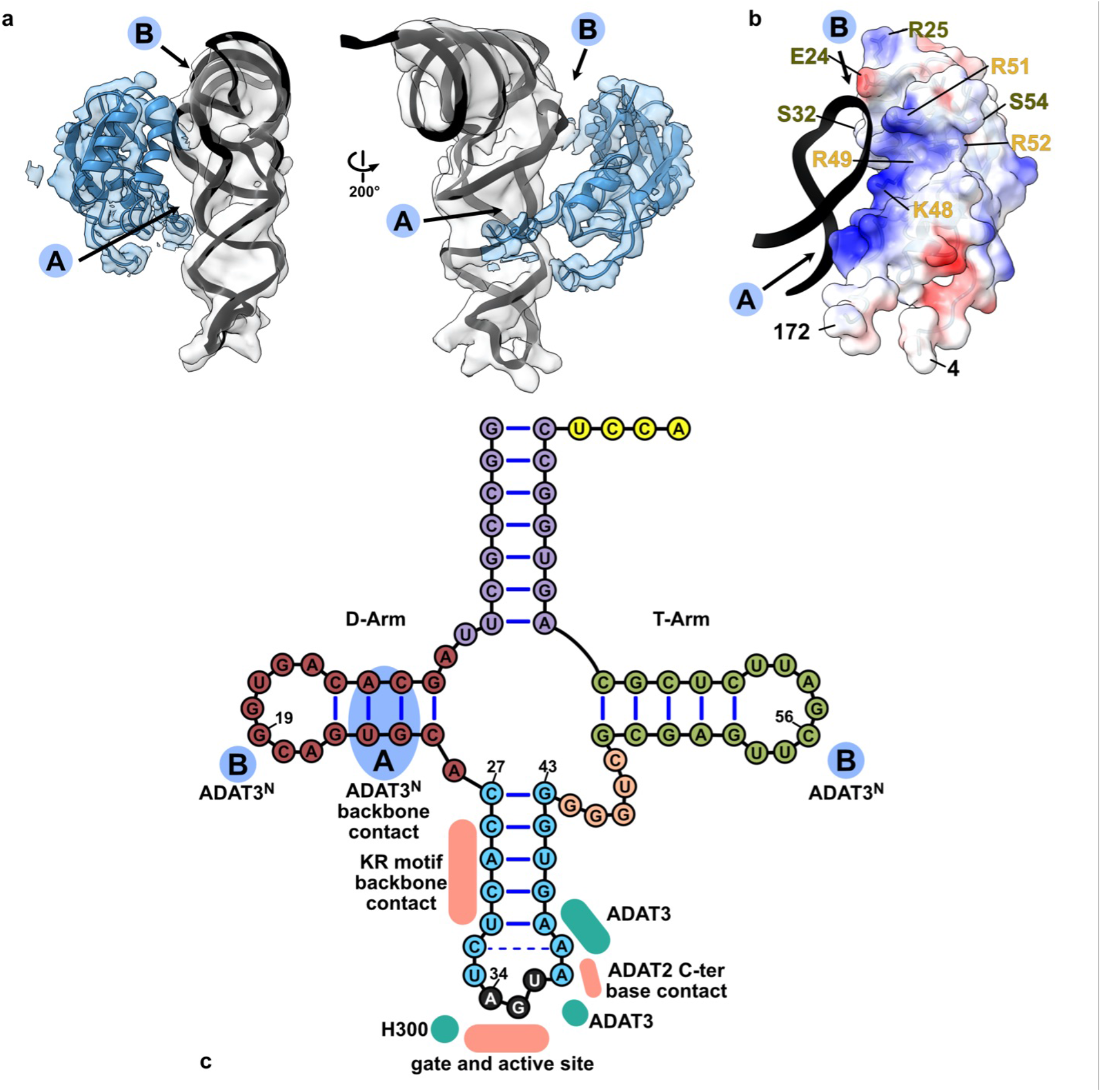
ADAT3 N terminal domain probes the 3-dimensional structure of the tRNA. (**a**) Two views of the ADAT3^N^ fitted in the post-processed EM map, in blue, with ‘a’ and ‘b’ regions as the two contact points of our model with tRNA, as indicated in C. (**b**) ADAT3^N^ including linkers (residues 4-172); surface model colored by vacuum electrostatic potential, positive charges in red and negative charges in blue, with D-Arm of tRNA in a black cartoon representation. Residues with a role in tRNA binding are annotated: from Ramos-Morales et al., 2021 in olive and from Liu et al., 2020 in yellow. (**c**) Schematic of the tRNA with interaction contacts. Anticodon stem loop interactions are represented as a summary of the ones indicated in figure 2D. The two ADAT3^N^ contact points, ‘A’ and ‘B’ are indicated in blue as above.

## Discussion

### Eukaryotic ADAT2/3 utilizes a multi-step tRNA recognition mechanism

The cryo-EM structure of the *Trypanosoma brucei* tRNA-bound ADAT2/3 complex pinpoints for the first time how the various RNA binding motifs of the eukaryotic deaminase interact with the full-length tRNA through a concerted mechanism. The structure, combined with available structural and biochemical data allows us to identify key conformational rearrangements that eukaryotic wobble base tRNA deaminases must undergo to correctly place the substrate in their active site for productive deamination. This allows us to propose the following improved model for a multi-step mechanism of tRNA recognition by ADAT2/3 (Fig. 6). In the free enzyme form, the N-terminal RNA binding domain of ADAT3 (ADAT3^N^) is flexibly linked to the deaminase core, and the C-terminus of ADAT2 (ADAT2^C^) containing the positively charged RNA-binding KR-motif is unfolded (Fig. 6a) (Liu et al., 2020; Ramos-Morales et al., 2021). In accordance with prior propositions, we suggest an initial binding step in which RNA is promiscuously captured via electrostatic interactions by these two RNA binding domains. Mutation or deletion of either the charged KR-motif of ADAT2^C^ (Ragone et al., 2011) or putative RNA binding residues in ADAT3^N^ (Ramos-Morales et al., 2021), lead to tRNA binding defects that point towards an initial and simultaneous RNA binding of both regions (Fig. 6b). Subsequently, we imagine that the anticodon loop is progressively directed into the deaminase active site, guided by the expansion of favorable protein-RNA contacts between the positively charged stretch along the linker region with the anticodon arm, leading to insertion of the anticodon loop into the catalytic pocket. Throughout the insertion, ADAT3^N^ and ADAT2^C^ might survey the integrity of the accurate tRNA structure determinants, with non-tRNA structures being rejected due to the lack of the correct contacts. Validated tRNAs can then enter through the molecular RY-gate, formed by cation-π interactions between the gate residue side chains, into the anticodon binding cleft of the joint ADAT2/3 deaminase core. Base stacking with the released gate residue tyrosine Y205^*Tb*ADAT3^ is enabled, allowing the nucleotides 33-36 to establish multiple interactions within the deep catalytic pocket. With the inserted anticodon loop, the unstructured portions in the ADAT2 C-terminus, notably the positively charged and essential KR-motif, adopt a folded conformation and help to correctly place the tRNA in a manner that is amenable for catalysis (Fig. 6c). For cognate tRNAs, the L-shape architecture, anticodon loop geometry, and the A_34_ nucleotide will perfectly align to the three main binding motifs in ADAT2/3, namely ADAT3^N^, ADAT2^C^ and joint deaminase core, to trigger the deamination reaction. We suggest that for substrate release, Brownian motions of each individual RNA binding site might initiate the discharge of non-cognate tRNAs (Fig. 6d). With hindsight, our structure may represent a transition state preceding product release with ADAT3^N^ and ADAT2^C^ showing hints of detached conformations, reflected in the reduced map resolution of both domains. We reason that when A_34_ is present, tRNA turnover is accelerated by the successful catalysis, leading to a fast product release. This is supported by the >1μM dissociation constant observed with an A_34_-containing tRNA in previous binding studies as compared to ∼180 nM observed with the C_34_-containing tRNA, a non-substrate which is inert to the reaction (Ragone et al., 2011). Conversely, non-A_34_-containing tRNAs, in the absence of catalysis, exhibit longer dwell times, leading to the conclusion that product release is driven by the energetics of the deamination reaction. Hence, a non-A_34_-containing tRNA may act as a competitive inhibitor for the reaction, which raises the question, how the cytoplasmic enzyme can be active *in vivo* in a pool of potential non-substrate inhibitors. Here, we can only assume that the effective concentration of free A_34_-tRNAs makes the difference, as non-substrate tRNAs may be sequestered from the available cytosolic pool by other tRNA targeting enzymes, like tRNA synthetases, translation factors and translating ribosomes. To conclude, we propose mechanistically, that for productive A_34_ deamination by eukaryotic ADAT2/3, all key enzyme-tRNA contacts, i.e. ADAT3^N^, ADAT2^C^ and the heterodimeric deaminase core, act in a concerted manner to capture the tRNA, probe for the correct geometry, and finally correctly position the substrate in the active site to trigger the deamination reaction.

**Fig. 6:**
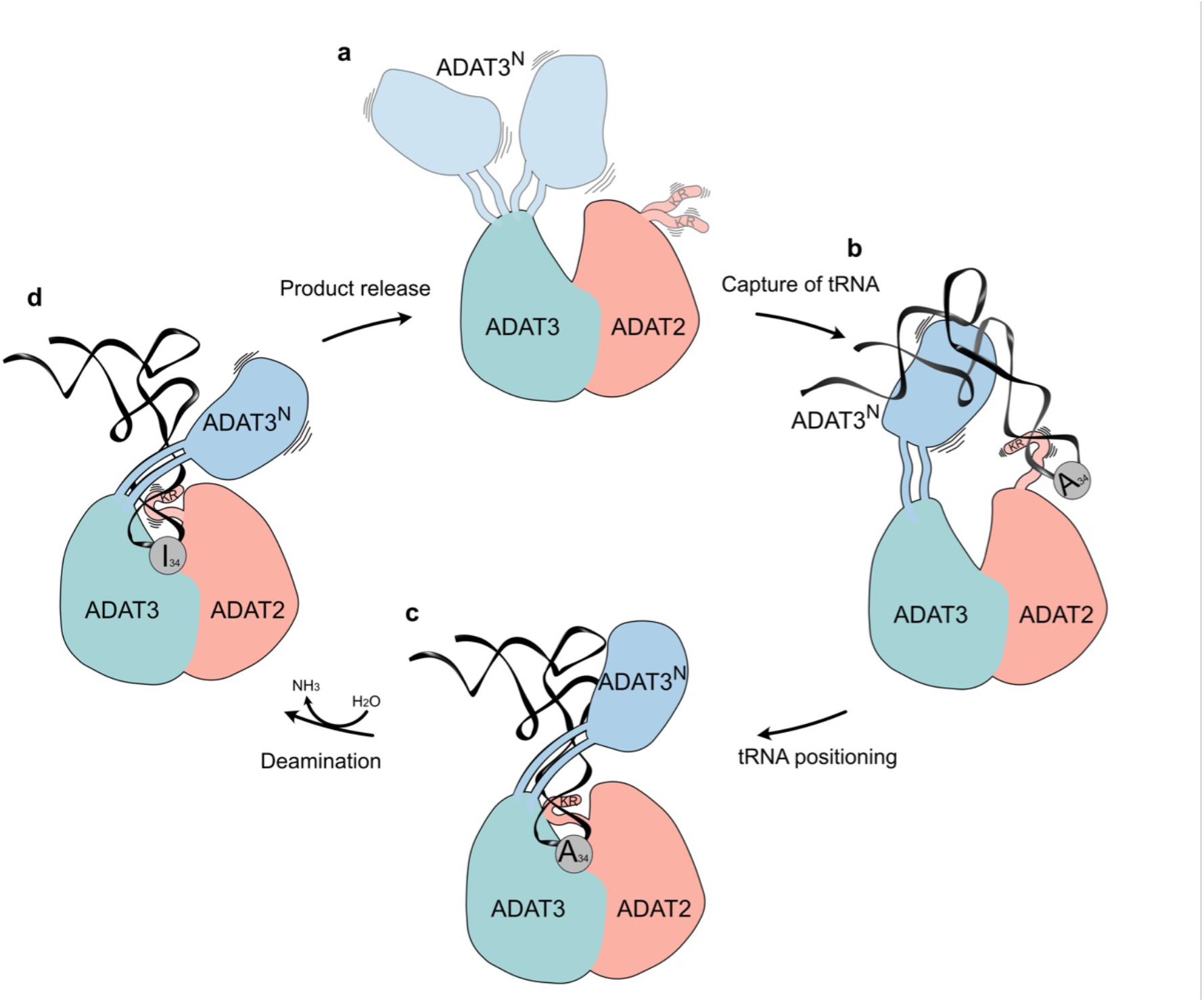
The hypothetical multi-step tRNA recognition mechanism of ADAT2/3. (**a**) **Ligand-free enzyme state:** in the free ADAT2/3 heterodimer ADAT2^C^ is disordered and ADAT3^N^ is flexibly linked. (**b**) **Initial capture:** the RNA recognition motifs in ADAT3^N^ and in ADAT2^C^ capture RNA; tRNAs are progressively guided by the extension of favorable interactions towards the anticodon binding cleft; hereby non-tRNA architectures will be rejected. (**c**) **Anticodon loop insertion into the active site:** The anticodon loop enters the active-site cleft through the RY-gate, and ADAT2^C^ folds into the major groove of the anticodon loop to position the ligand for the reaction. (**d**) **Substrate release:** Detachment of the individual RNA binding motifs initiates tRNA release.

### Asymmetry of the ADAT2/3 heterodimer led to diversification of its tRNA targets

Several structures of the bacterial deaminase TadA, combined with biochemical analysis suggested that the bacterial enzyme makes sequence-specific contacts in their active site to efficiently target its single substrate tRNA^Arg^_ACG_. It has been proposed that substrate diversification to multiple tRNAs by ADAT2/3 was the result of active site mutations that effectively relaxed the need for sequence-specific recognition (Elias and Huang, 2005). A feasible evolutionary scenario for increased substrate diversity by the eukaryotic enzymes may have involved a gene-duplication of the ancestral ADAT2 protein to yield ADAT3 and the enzyme acquired mutations that lead to active-site relaxation (Auxilien et al., 1996; Elias and Huang, 2005; Gerber and Keller, 1999; Rubio et al., 2007; Torres et al., 2021; Zhou et al., 2014). In this intermediate state, ADAT2 was still likely able to catalyze the reaction in a single substrate á la TadA. However, appearance of the ADAT3 subunit, and its ability to heterodimerize, allowed it to abandon sequence-specific recognition and acquire additional sequence-independent RNA binding elements located distant from the catalytic center, reducing the importance of the active-site in tRNA recognition and finally leading to a multi-step binding mechanism. We observe that while the bacterial TadA homodimer almost exclusively interacts with the anticodon loop bases C_32_-A_38_, the additional eukaryotic domains add manifold sequence-nonspecific contacts along the anticodon arm up to the elbow region of the tRNA (Fig. 5C and Extended Data Fig. 5). As a relic of its origin, in extant ADAT2/3 enzymes, the ADAT2 enzyme is still able to form homodimers *in vitro*, yet these are inactive (Ramos-Morales et al., 2021; Spears et al., 2011b; PDB: 3DH1), reflecting the evolved dependency on ADAT3 for enzyme activity.

The tRNA-bound cryo-EM structure of the ADAT2/3 heterodimer presented here allows us to unambiguously pinpoint the exact function of all crucial tRNA binding features that differentiate ADAT2 and ADAT3. ADAT2 specific features are the RY-gate, necessary for access to the deaminase active site and the essential C-terminal KR-motif, which, intrinsically unfolded in the ligand-free state, contributes to the correct placement of the anticodon stem loop once the tRNA is bound. ADAT3, on the other hand, has acquired an indispensable additional N-terminal RNA binding domain to probe global tRNA geometry through extended sequence-independent contacts to the tRNA phosphodiester backbone; a somewhat similar case has been observed for the tRNA (guanine(37)-N1)-methyltransferase Trm5, that also recognizes the full tRNA L-shape via its “sensor” D1 domain, which probes for the intact 19-56 base pair before recruiting the “effector” catalytic domain (Christian and Hou, 2007; Goto-Ito et al., 2017; Wang et al., 2017). In a similar manner, also the tRNA anticodon modifying enzymes TilS and TiaS recognize features of their cognate tRNA via domains which are distal from the active center, albeit base specific in these cases (Numata, 2015). Interestingly, while the evolutionarily conserved A_34_ pocket in ADAT2/3 appears of relatively rigid character, the distal eukaryote-acquired RNA binding features show a certain flexibility either via connection through linkers (ADAT3^N^) or induced-fit adaptation of the protein portion (ADAT2^C^). This flexibility is most likely the key to the accommodation of multiple tRNAs with divergent anticodon loop sequences (Extended Data Fig. 5). Furthermore, we find it interesting that ADAT3s of different eukaryotic clades all have evolved different loops to obstruct their pseudo-active site, potentially to prevent tRNAs to bind in an unproductive manner (Extended Data Fig. 6) (Liu et al., 2020; Ramos-Morales et al., 2021).

## Conclusion

The cryo-EM structure of the full-length ADAT2/3 heterodimer bound to a tRNA delivers the long-sought rationale of how a sequence specific single-substrate wobble base deaminase has evolved into a multi-tRNA acceptor. Key to its evolution were the acquisition of additional RNA binding domains and motifs (ADAT3^N^ and ADAT2^C^), and the development of a multi-step tRNA recognition mechanism, which was only possible after gene duplication allowing the two subunits to diverge and undergo neo-functionalization. The acquired distal RNA-binding motifs provide additional ductile affinity to the ligand through induced-fit mechanisms, but moreover they serve to authenticate tRNAs via their global 3D features; delicately counterbalancing multi-sequence recognition while keeping the enzyme’s mutagenic potential in check.

## Material and Methods

### Protein Expression and Purification

For structure determination full-length ADAT2 (*Tb*927.8.4180, with two point mutation that improve solubility: A87G/C117S) and full-length His-tagged ADAT3 (*Tb*927.11.15280) were co-expressed in Hi5 cells using the multibac and Bigbac insect cell expression systems (Fitzgerald et al., 2006; Weissmann et al., 2016). Cells were resuspended in lysis buffer (20 mM Tris pH 7.5; 200 mM NaCl; 20 mM Imidazole; 2% Glycerol; supplemented with 1 mM of PMSF, 2 µg/mL of DNase and 25 µg/mL of RNase) and lysed by sonication (5 minutes, 30% Amplitude, 5 seconds on, 5 seconds off; Vibra-cells, Sonics). The lysate was clarified by centrifugation (40,000 g, 30 minutes) and loaded into a 5 ml HisTrap HP affinity chromatography column (Cytiva), that was washed with 40 column volumes of high salt buffer (20 mM Tris pH 7.5; 1 M KCl; 50 mM Imidazole) and 40 column volumes of low salt buffer (20 mM Tris pH 7.5; 200 mM NaCl; 50 mM Imidazole). The sample was eluted with elution buffer (20 mM Tris pH 7.5; 200 mM NaCl; 600 mM Imidazole) and flushed through a HiTrap Q HP 5 mL anion exchange chromatography column (Cytiva) to remove charged impurities. After 3C protease tag cleavage, the sample was concentrated through centrifugation in Centricon filters and finally purified through size exclusion chromatography on a HiLoad 16/600 Superdex 200 column (Cytiva) pre-equilibrated with SEC buffer (20 mM Hepes pH 7.5; 100 mM NaCl; 2 mM DTT). All steps were performed at 4°C.

For biochemical assays, *Tb*ADAT2/3 was recombinantly expressed in *E. coli* as previously described. Briefly, the coding sequences of *Tb*ADAT2 and *Tb*ADAT3 were cloned into expression vectors pETDuet-1 and pET-28(a)+ and used to transform *E. coli* BL21-IRL strain. For expression, a 10 mL overnight culture was added to 1.5 L of prewarmed 2XYT media and grown at 37°C to an OD600 of 0.6–0.8. Recombinant protein expression was then induced with isopropyl β-d-1-thiogalactopyranoside (IPTG, 0.5 mM final concentration) overnight at 25°C. The following procedures for protein purification were carried out at 4°C. Cells were pelleted, suspended in lysis buffer (20 mM Tris pH 8.0, 1 M NaCl, 5 mM β-mercaptoethanol, 20 mM Imidazole, protease inhibitor cocktail and 0.1% NP40) and lysed by sonication with a Branson Sonifier 450 (three times, 30-s intervals, 30-s rest between sonication). The resulting lysate was clarified by centrifugation at 40,000g for 30 min followed by a second centrifugation step at 55,000g for 30min. Clear lysate was collected and loaded onto a 1mL HisTrap HP column (Cytiva, Ni^2+^-nitrilotriacetic acid column). The column was washed with the lysis buffer followed by a lower salt wash containing 20 mM Tris pH 8.0, 100 mM NaCl, 5 mM β-mercaptoethanol, and 50 mM Imidazole. The bound protein was eluted with 600 mM Imidazole. Peak fractions were pooled then passed through a PD-10 desalting column (Cytiva) and stored as aliquots at -80°C in buffer containing 100 mM Tris pH 8.0, 100 mM NaCl, 0.5 mM MgCl_2_, 0.2 mM EDTA, 2mM 1,4-dithiothreitol, and 15-20% glycerol.

### Thermostability assay

Thermal stability of *Tb*ADAT2/3 was determined by differential scanning fluorimetry (DSF). Briefly, *Tb*ADAT2/3 (in a final concentration of 4 µM) was mixed with SYPRO Orange in DSF buffer (20 mM Hepes pH 8; 200 mM NaCl; 2 mM DTT) and incubated in a temperature gradient for denaturation. The melting temperature was approximated to the midpoint between maximum and baseline signals of the SYPRO Orange fluorescence (Mariaule et al., 2014).

### ADAT2/3 ligand sequencing

Nucleic acids co-eluted in the size-exclusion peak-fraction of recombinantly expressed *Tb*ADAT2/3 derived from *E. coli* as indicated by an absorption 260/280 ratio of approximately 1.7. Nucleic acids were phenol-chloroform extracted and precipitated with sodium acetate. The sequencing reaction was prepared following a tRNA sequencing protocol with the NEXTflexTM Small RNA-Seq Kit v3 (Zheng et al., 2015). Adapter removal, 5’ and 3’ trimming of the result sequences and removal of sequences <15 bases resulted in 3577188 reads which were aligned with bowtie2 to an *E. coli* or *E. coli* tRNA index, respectively, and resulted in a 57.69% or 25.96%, respectively, overall alignment rate.

### tRNA synthesis for cryo-EM

The *Trypanosoma brucei* threonine tRNA (*Tb*tRNA^Thr^_CGU_) sequence was cloned in a pUC19 vector, between a T7 promoter and a BstNI cleavage site. The vector was linearized with BstNI and used as a template for in vitro transcription. The reaction was carried out in T7 reaction buffer (40 mM Tris pH 8; 5 mM DTT; 1 mM spermidine; 0.01% Triton X-100; 30 mM MgCl2), with 4 mM of each NTP. For a 10 mL reaction, 0.5 mg of linear DNA and 0.4 mg of T7 RNA polymerase were used. The reaction was incubated at 37 °C for 8 hours. The transcribed RNA was isopropanol precipitated, Urea-PAGE purified and desalted following standard procedures. The tRNA^Thr^ was refolded by heating at 95°C for 10 minutes in 10 mM Tris pH 7.5 and 50 mM NaCl, slow cool down and adding MgCl_2_ to 1 mM when it reached 37 °C. After refolding, *Tb*tRNA^Thr^_CGT_ was further purified by size exclusion chromatography, in a Superdex 200 Increase 10/300 GL size exclusion chromatography column (Cytiva) pre-equilibrated with refolding buffer (10 mM Tris pH 7.5 and 50 mM NaCl, 2 mM MgCl^2^).

### Cryo-EM sample preparation

For complex formation, ADAT2/3 heterodimer and tRNA were mixed and diluted in interaction buffer (20 mM Hepes pH 7.5; 50 mM NaCl; 2 mM MgCl2; 0.5 mM TCEP) to a final concentration of 32 µM and 35 µM, respectively. Excess tRNA was removed through size exclusion chromatography on a Superdex 200 Increase 3.2/300 column (Cytiva). UltrAuFoil R 1.2/1.3 300 Au mesh (Quantifoil) grids were glow discharged with residual air for 30 seconds, on each side in a PELCO easiGlow device operated at 30 mA. 2 µL of sample were applied in each side of the grid before blotting for 3 seconds (blot force 0) and vitrified in liquid ethane using a Vitrobot MARK IV (Thermo Fisher Scientific) operated at 4 °C and 100% humidity.

### Cryo-EM Data acquisition

Micrographs were collected with a 300kV Titan Krios (Thermo Fisher Scientific) electron microscope, equipped with a K2 Summit direct electron detector and a GIF Quantum energy filter (Gatan). Data were acquired using serialEM (Mastronarde, 2005) at a nominal magnification of 165,000, resulting in a pixel size of 0.81 Å. For a better sampling of particle orientations, the stage was tilted 30°; to reduce beam induced motion, the beam size was increased to 650 nm. Movies were acquired for 10 seconds at a dose of 3 e/pixel/s, resulting in a total dose of 55.00 e/A^2^ at the sample level, fractionated into 80 movie frames. Three movies were acquired per hole, for a total of 6,145 movies, with a defocus range from -1.0 to -1.8 µm.

### Cryo-EM Data processing

Movies were motion corrected (using a 5×5 patch model) and patch CTF estimated (3×3 patch model) in Warp. A total of 1,156,506 particles were picked and extracted (300 pixel box) from the corrected micrographs using the BoxNet2Mask_21080918 model (Tegunov and Cramer, 2019). After initial 2D classification in CryoSPARC (Punjani et al., 2017), we could identify three main particle subsets: tRNA, with 62,505 particles; ADAT2/3 heterodimer, with 83,218 particles; and ADAT2/3-tRNA complex, with 559,713 particles. One round of 3D classification generated three ab-initio reconstructions followed by hetero refinement, from which the main class, with 427,165 particles were 2D classified. Blurred 2D classes were discarded and the remaining 379,795 particles were once more 3D classified by generating 4 ab-initio reconstructions followed by hetero refinement. The 105,718 particles from the best class were locally CTF refined, and further refined in a non-uniform refinement to generate the final 3.6A map that was post-processed using DeepEMhancer (Sanchez-Garcia et al., 2021).

### Model generation with AlphaFold

The model used to build the ADAT2/3 structure in the EM map was generated with AlphaFold. The sequences of ADAT3 and ADAT2, connected via a 45-residue AGS linker, were submitted to the AlphaFold colab notebook, and the 5 resulting models were compared based on their predicted IDDT and PAE.

(https://colab.research.google.com/github/sokrypton/ColabFold/blob/main/AlphaFold2.ipynb#scrollTo=G4yBrceuFbf3)

### Model building

The final map was sharpened or blurred in Coot 0.9.4 (Casañal et al., 2020) for easier interpretation. For the model building, the deaminase domain of ADAT2 and ADAT3 were built based on a model of *Tb*ADAT3 calculated by Phyre^2^ (Kelley et al., 2015) using the *Mm*ADAT3 (PDB: 7NZ7) (Ramos-Morales et al., 2021) as a template, and an AlphaFold generated model of *Tb*ADAT2/3. The AlphaFold generated N-terminal domain of ADAT3 was jiggle-fitted in the DeepEMhancer post-processed map using the traced linkers as extra restraints. For the tRNA, we rigid-body fitted the crystal structure of yeast phenylalanine tRNA (PDB: 1EHZ) (Shi and Moore, 2000) into the density map, mutate nucleotides to correspond to the *Tb*tRNA^Thr^_CGU_ sequence, and refined only the portion relative to the anticodon stem loop bases 28-42. The model was then refined against the final map in an interactive manner with Ramachandran, secondary structure, base pair and base stacking restraints, using both Coot 0.9.4 and Phenix (Liebschner et al., 2019). Alignments were made with pymol or clustal, and represented with ESPript 3 (Robert and Gouet, 2014; Thompson et al., 1994).

### Deamination assay

To prepare substrate for deamination assays, full-length tRNA^Thr^_AGU_ was in vitro transcribed using a T7 promoter and internally labeled with [α-^32^P]-ATP as previously described (Rubio et al., 2007, 2017). Single turnover kinetics were performed in reaction buffer containing 40 mM Tris pH 8, 5 mM MgCl_2_, and 10 mM DTT. Enzyme was added to the reaction in excess to labeled tRNA substrate (1nM) and incubated at 27°C as previously described (Ragone et al., 2011). Aliquots from the reaction were taken at various time points up to one hour (with exception of R159Y^*Tb*ADAT2^, which was up to four hours). Mutants with no detectable activity were incubated for 24 h. Samples were quenched via phenol extraction followed by ethanol precipitation. The pelleted samples were then resuspended and treated with Nuclease P1 overnight (18 h) at 37°C. Digested samples were then dried with high heat in a SpeedVac DNA 110 concentrator system (Savant). Dried sample pellets were resuspended in water and spotted onto a cellulose thin-layer chromatography (TLC) plate (Merck). Products were resolved for 2 h in one dimension in solvent C containing 0.1 M sodium phosphate (pH 6.8): ammonium sulfate: n-propanol (100:60:2 [v/w/v]). The TLC was dried and exposed to a PhosphorImager Screen overnight. Results were visualized using Typhoon FLA 9000 (GE) and analyzed using ImageQuant software. Fraction of inosine formed was calculated using pI/(pA + pI). The fraction A_34_ to I conversion was plotted against time and fit using MATLAB software to a single exponential curve [f = a(1 − e ^−kt^)], where f represents product formed, a denotes product formed at the end point of the reaction, k signifies k_obs,_ and t is time.

## Data availability

EM maps are available at EMDB: XXXX and the model is deposited at PDB: XXXX

## Acknowledgements

We acknowledge Martin Pelosse for support in using the Eukaryotic Expression Facility at EMBL Grenoble, and Sarah Schneider for support in using the EM Facility at EMBL Grenoble. This work benefited from access to the cryo-EM platform of the Structural and Computational Biology Unit at EMBL Heidelberg. We acknowledge the HTX Team for the thermostability assay. We acknowledge the EMBL Genomic Core Facility for the tRNA sequencing. This work used the platforms of the Grenoble Instruct-ERIC center (ISBG; UAR 3518 CNRS-CEA-UGA-EMBL) within the Grenoble Partnership for Structural Biology (PSB), supported by FRISBI (ANR-10-INBS-0005-02) and GRAL, financed within the University Grenoble Alpes graduate school (Écoles Universitaires de Recherche) CBH-EUR-GS (ANR-17-EURE-0003). We thank Aline Le Roy and Christine Ebel, for assistance and access to the Protein Analysis On Line (PAOL) platform. The authors thank the Kowalinski lab members for discussion and comments throughout the course of the project. We also thank Andrew McCarthy and Sebastian Falk for comments on the manuscript. The present work was funded in part by the National Institutes of Health Grant GM084065-11 to J.D.A. The work was supported by a grant from the French Agence Nationale de la Recherche to E.K. (ANR-20-CE11-0016).

## Author contributions

EK designed the study. EK, LGD, LT conducted experimental work. FW collected the tilted cryo-EM data. LGD processed the cryo-EM data and built the model. LGD and EK interpreted the data and wrote the manuscript. MAR performed experiments and interpreted the data. JDA interpreted the data and wrote and edited the manuscript. AZ performed experiments and interpreted the data.

## Notes

### Competing Interest Statement

The authors have declared no competing interest.

### Summary of Updates

Orchid IDs included, change of licence

