## Extended_Data for "Sequence-independent substrate selection by the eukaryotic wobble base deaminase ADAT2/3 involves multiple protein domains and distortion of the tRNA anticodon loop"

**Extended Data Fig. 1: Biochemical characterization of *Tb*ADAT2/3.**
**(a)** SDS-PAGE of fractions of the purified ADAT2 (24 kDa) and ADAT3 (43 kDa) complex. **(b)** Thermostability assay of the ADAT2/3 heterodimer, with a melting temperature of 53.5 ˚C. **(c)** Different RNAs identified to co-purify with ADAT2/3 from overexpression in *E.coli*. The first graph represents the high abundant tRNA in comparison with all other RNA hits. The second represents the tRNA abundances compared to all tRNA identified, with $tRNA^{Thr}$ and $tRNA^{Pro}$ as the most abundant. **(d)** SEC-MALLS experiment with different molecular ratios of added tRNA, indicating a complex stoichiometry of 1:1:1 ADAT2:ADAT3:tRNA. **(e)** Model of ADAT2/3 as predicted by AlphaFold in cartoon representation. ADAT2 in salmon, ADAT3 $^{CDA}$ in teal and $ADAT3^{N}$ in blue. Regions with IDDT lower than 50 were hidden from the representation. **(f)** Statistics from the AlphaFold model presented in E.

**Extended Data Fig. 2: Structure determination of *Tb*ADAT2/3 bound to tRNA by CryoEM.**
**(a)** Example of a representative micrograph (resulting from the averaging of the motion corrected frames of a movie). **(b)** Patch CTF estimation from warp, where we see the relative tilt of the stage generating a defocus gradient in the micrograph. A more precise initial CTF estimation is of particularly importance in the case of tilted data collection. **(c)** First round of 2D classification used to separate the tRNA bound ADAT2/3 particles from the free tRNA or free ADAT2/3 particles. **(d)** First round of 3D classification to further remove free tRNA or free ADAT2/3 particles from our tRNA bound ADAT2/3 dataset. **(e)** Second round of 2D classification, to further remove low resolution particles. **(f)** Second round of 3D classification, to increase particle homogeneity. **(g)** Final refinement step, combining local CTF refinement with non-uniform refinement. **(h)** Post processing of the final refinement, performed by DeepEMhancer, that increased the connectivity of the map and allowed for the interpretation of the $ADAT3^{N}$ (colored as in figure 1, with ADAT2 in salmon, $ADAT3^{N}$ in blue, $ADAT3^{CDA}$ in teal and the tRNA in black). **(i)** Local resolution map calculated by CryoSPARC, displayed in Chimera, with level 0.1. **(j)** Final map GSFSC (Gold Standard Fourier Shell Correlation) as calculated by cryoSPARC indicates a resolution of 3.62A. **(k)** Angular distribution map of particle orientation as calculated by cryoSPARC.

**Extended Data Fig. 3: Anticodon loop, active site and molecular RY gate.**
**(a)** Two views of the anticodon loop region of the tRNA, with the EM density represented as a blue mesh, indicating that all bases are well resolved. Hydrogen bond of the non-canonical base pair between $C_{32}$ and $A_{38}$ represented in cyan. **(b)** Active site with the EM density represented as a blue mesh. No density for the catalytic glutamate E92 side chain and for the zinc coordinating water molecule can be observed. **(c)** Comparison with the catalytic site of the yeast crystal structure (PDB: 7BV5). Water molecule in red, zinc atom in yellow, dotted lines representing the zinc coordination, side chains colored by heteroatom. **(d)** Gate residues R159 and Y205 of ADAT2. The absence of density for the side chain of R159 suggests its flexibility. **(e)** Internalization of base $U_{36}$ that is sandwiched in a π-π stacking interaction with base $C_{32}$ and gate residue Y205. Hydrogen bond of the non-canonical base pair between $C_{32}$ and $A_{38}$ in cyan. **(f)** Detail of the closed conformation of the molecular RY gate as show in the structure of *Mm*ADAT2/3 (PDB: 7NZ7). Electron density map contoured at 1 σ is shown as blue mesh. **(g & h & i)** Comparison of an ADAT3 conserved loop between eukaryotic bound and unbound structures: **(g)** *Tb*ADAT2/3; **(h)** *Mm*ADAT2/3 (PDB: 7NZ7), with overlay of the tRNA from the *Tb*ADAT2/3 structure for visualization purposes; **(i)** bacterial RNA bound (*Sa*TadA, PDB: 2B3J). All RNA in black, and side chain and nucleosides colored by heteroatom. **(j & k)** Single turnover assays of wild-type *Tb*ADAT2/3 and selected *Tb*ADAT2 gate mutants co-expressed with wild-type *Tb*ADAT3. $A_{34}$ to $I_{34}$ conversion was measured over 1 h in the presence of excess enzyme; except for $R159Y^{TbADAT2}$, which was measured over 4 h. The dashed line in K represents the maximum A-to-I fraction in the wild-type assay. The fraction of inosine formed was plotted against time and fit to a single

exponential curve [$f = a(1 - e^{-kt})$], where f represents product formed, a denotes product formed at the end point of the reaction, k signifies $k_{obs}$, and t is time.

**Extended Data Fig. 4: The N-terminal domain of ADAT3 is involved in tRNA recognition and binding.**

**(a)** Alignment of ADAT3 N-terminal domains, with the sequences for *trypanosoma brucei*, human, mouse and yeast. Structural elements predicted by AlphaFold for *Tb*ADAT3$^N$ are annotated with alpha-helices as cylinders and beta-sheets as arrows. Residues identified as important for tRNA binding or deamination in the corresponding species are annotated as olive triangles above for *Mm*ADAT3$^N$ and yellow triangles below for *Sc*ADAT3$^N$.The annotated blue box indicates a kinetoplast specific region, which is predicted to be disordered. Other colors and boxes in the alignment represent relative conservation above 70%. **(b)** CryoSPARC non-uniform refinement map of ADAT2/3, with the tRNA as cartoon, colored as in figure 1, the blurred density accounting for the ADAT3$^N$. **(c)** Same as B, but EM density map after post-processing with DeepEMhancer, that improved the map quality in the deaminase core, and allowed to jiggle-fit the ADAT3$^N$ model.

**Extended Data Fig. 5: Comparison between *Sa*TadA and *Tb*ADAT2/3 interaction with tRNA.**

**(a & b)** Schematic overview of the anticodon stem loop region of the tRNA and its interactions with **(a)** *Sa*TadA and **(b)** *Tb*ADAT2/3, respectively. Zinc coordinating sites are highlighted in yellow. ADAT2 residues are represented in salmon, ADAT3 residues in teal. Residues of the TadA α-protomer of the homodimer (corresponding to ADAT2 relative to the ligand) in grey and residues of the β-protomer (corresponding to the ADAT3 relative to the ligand) in pink. **(c & d)** Schematic of the whole tRNA with interaction contacts for *Sa*TadA and *Tb*ADAT2/3. Anti-codon stem loop contacts are represented as a summary of the ones in A and B**.** Since no structure of TadA bound to full length tRNA is available, the same schematic tRNA is used for both. Colors as in A and B.

**Extended Data Fig. 6: ADAT3 pseudo-active site plug.**

**(a, b, c & d)** Comparison of the pseudo active sites plug in **(a)** *Tb*ADAT3 (with coulomb density as blue mesh), **(b)** *Mm*ADAT3 (PDB: 7NZ7), and **(c)** *Sc*ADAT3 (PDB:7BV5). **(d)** *Tb*ADAT2 active site as a comparison. Zinc atom in yellow, dotted lines representing the zinc coordination, side chains colored by heteroatom. All ADAT3 sites are obstructed by ADAT3 protein loops ("plugs"). **(e & f)** Sequence alignment of kinetoplast, mammalian and yeast ADAT3s regions representing the "plug" protein loops. The residues in yellow boxes correspond to the plug in each eukaryotic clade. A gray star represents the reaction inert residue that substitutes the glutamic acid in the active site. The plug is present in clade specific loops or non-conserved regions, indicating that this feature has divergently evolved. Colors and boxes in the alignment represent relative conservation above 70%. **(g)** Expression and purification test of wild-type *Tb*ADAT2 / wild-type *Tb*ADAT3 in comparison with wild-type *Tb*ADAT2 / *Tb*ADAT3 Y236 mutants. All tested point mutations lead to insoluble protein reaffirming the importance of Y236 as acquired additional $Zn^{++}$ coordinating residue. Total fraction after lysis of the insect cells; soluble fraction after the lysate clarification; elution from the Ni-NTA resin capture.

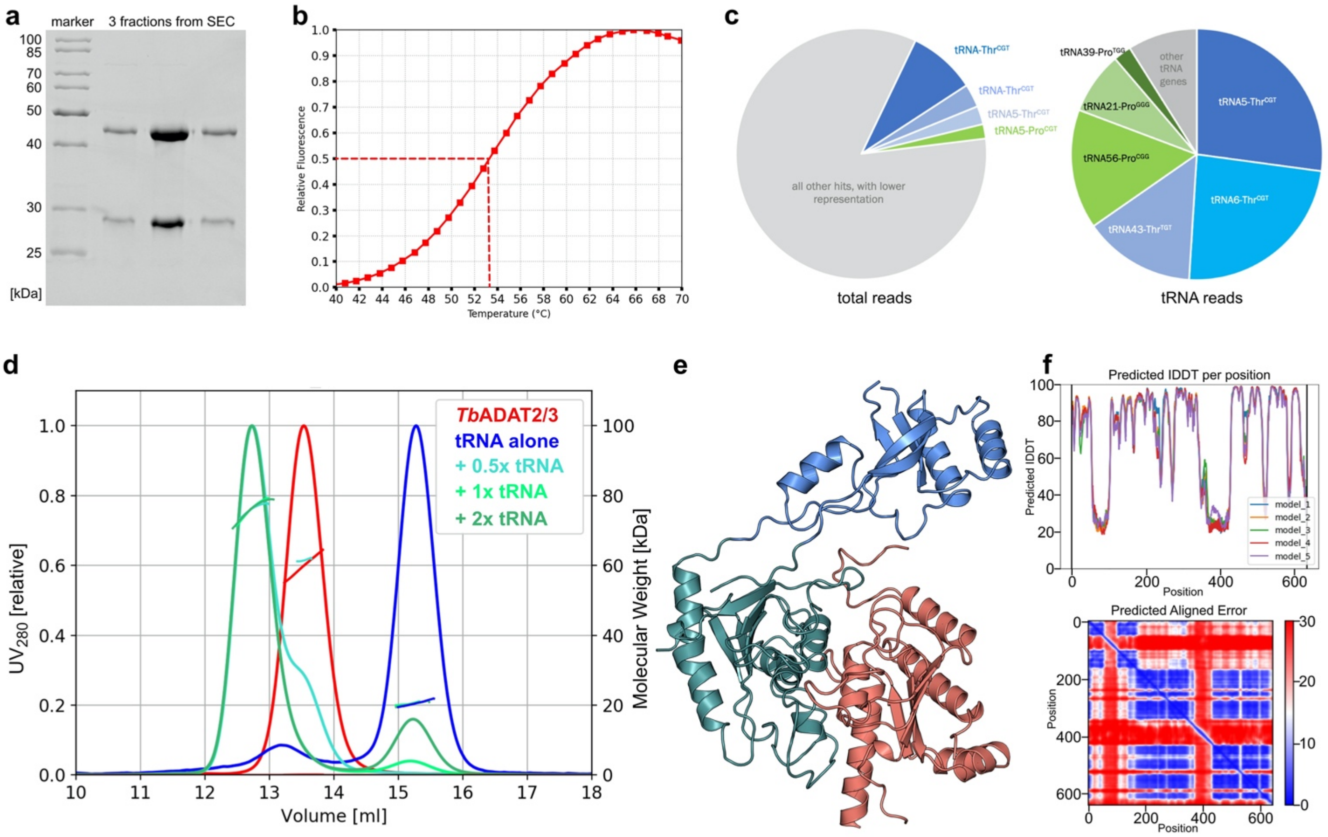
a
marker
3 fractions from SEC
100
85
70
60
50
40
30
25
[kDa]
b
Relative Fluorescence
Temperature (°C)
c
tRNA-Thr^CGT
tRNA-Thr^CGT
tRNA5-Thr^CGT
tRNA5-Pro^CGT
all other hits, with lower representation
total reads
tRNA39-Pro^TGG
other tRNA genes
tRNA21-Pro^GGG
tRNA56-Pro^CGG
tRNA43-Thr^TGT
tRNA5-Thr^CGT
tRNA6-Thr^CGT
tRNA reads
d
TbADAT2/3
tRNA alone
+ 0.5x tRNA
+ 1x tRNA
+ 2x tRNA
UV280 [relative]
Molecular Weight [kDa]
Volume [ml]
e
f
Predicted lDDT per position
Predicted lDDT
Position
model_1
model_2
model_3
model_4
model_5
Predicted Aligned Error
Position
Position

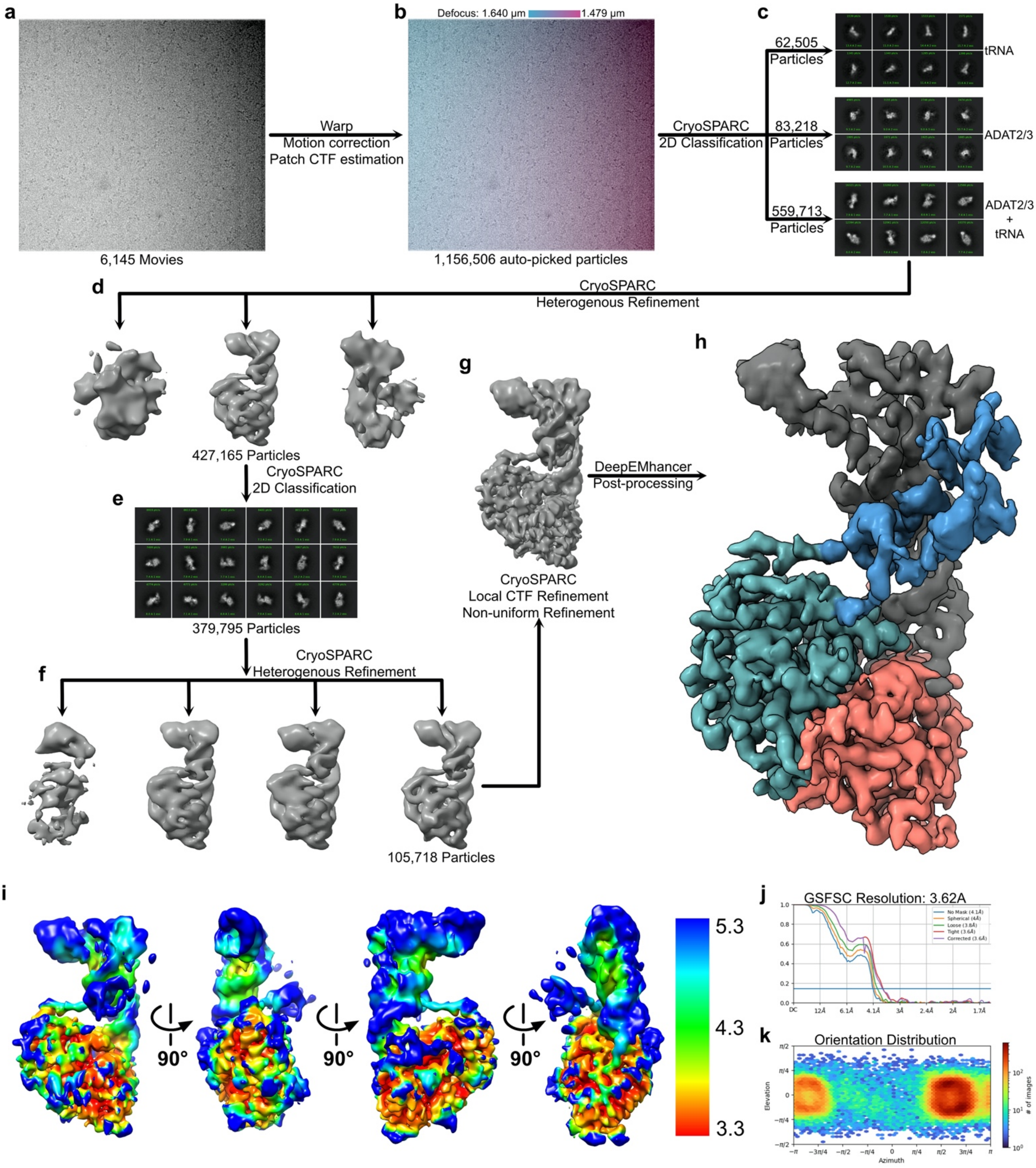
a
b
Defocus: 1.640 μm
1.479 μm
c
62,505
Particles
tRNA
Warp
Motion correction
Patch CTF estimation
CryoSPARC
2D Classification
83,218
Particles
ADAT2/3
559,713
Particles
ADAT2/3
+
tRNA
6,145 Movies
1,156,506 auto-picked particles
d
CryoSPARC
Heterogenous Refinement
h
g
427,165 Particles
CryoSPARC
2D Classification
DeepEMhancer
Post-processing
e
379,795 Particles
CryoSPARC
Local CTF Refinement
Non-uniform Refinement
CryoSPARC
Heterogenous Refinement
f
105,718 Particles
i
90°
90°
90°
5.3
4.3
3.3
j
GSFSC Resolution: 3.62A
k
Orientation Distribution
Elevation
Azimuth
### of images

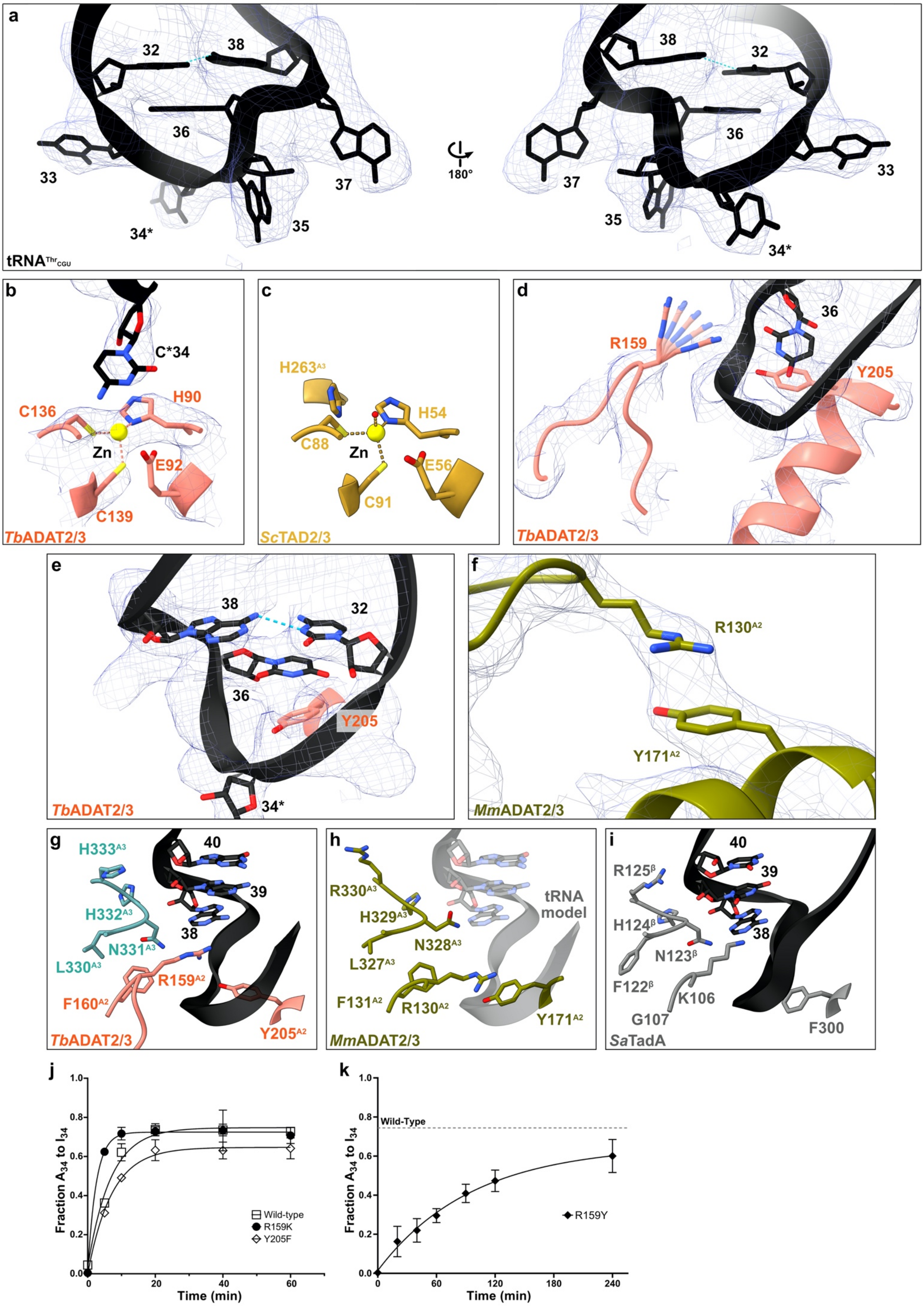

a
32
38
36
33
34*
35
37
180°
tRNAThrCGU
b
C*34
C136
H90
Zn
E92
C139
TbADAT2/3
c
H263A3
H54
C88
Zn
E56
C91
ScTAD2/3
d
R159
36
Y205
TbADAT2/3
e
38
32
36
Y205
34*
TbADAT2/3
f
R130A2
Y171A2
MmADAT2/3
g
H333A3
40
H332A3
39
38
N331A3
L330A3
R159A2
F160A2
Y205A2
TbADAT2/3
h
R330A3
H329A3
N328A3
L327A3
F131A2
R130A2
Y171A2
tRNA model
MmADAT2/3
i
40
R125β
39
H124β
38
N123β
F122β
K106
G107
F300
SaTadA
j
Fraction A34 to I34
Time (min)
Wild-type
R159K
Y205F
k
Wild-Type
Fraction A34 to I34
Time (min)
R159Y

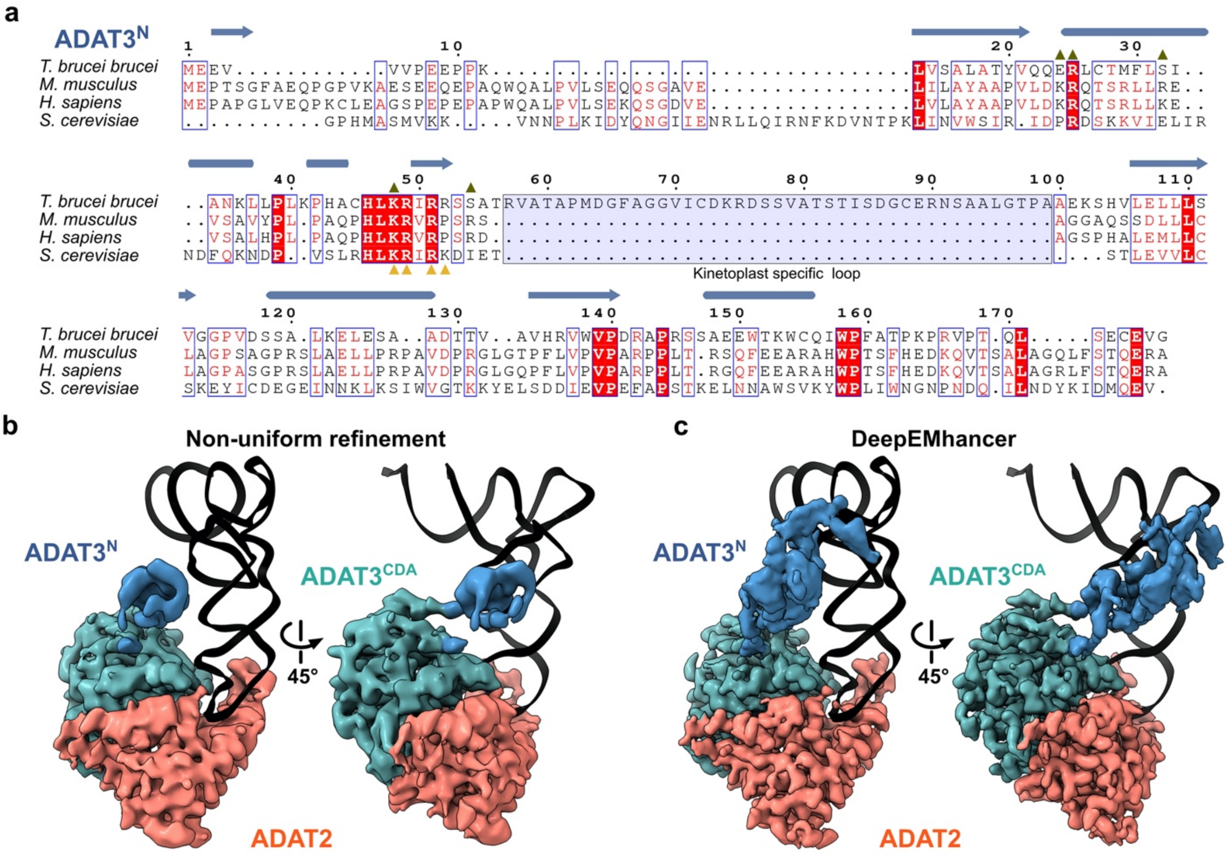

a
ADAT3N
T. brucei brucei
M. musculus
H. sapiens
S. cerevisiae
Kinetoplast specific loop
b
Non-uniform refinement
ADAT3N
ADAT3CDA
45°
ADAT2
c
DeepEMhancer
ADAT3N
ADAT3CDA
45°
ADAT2

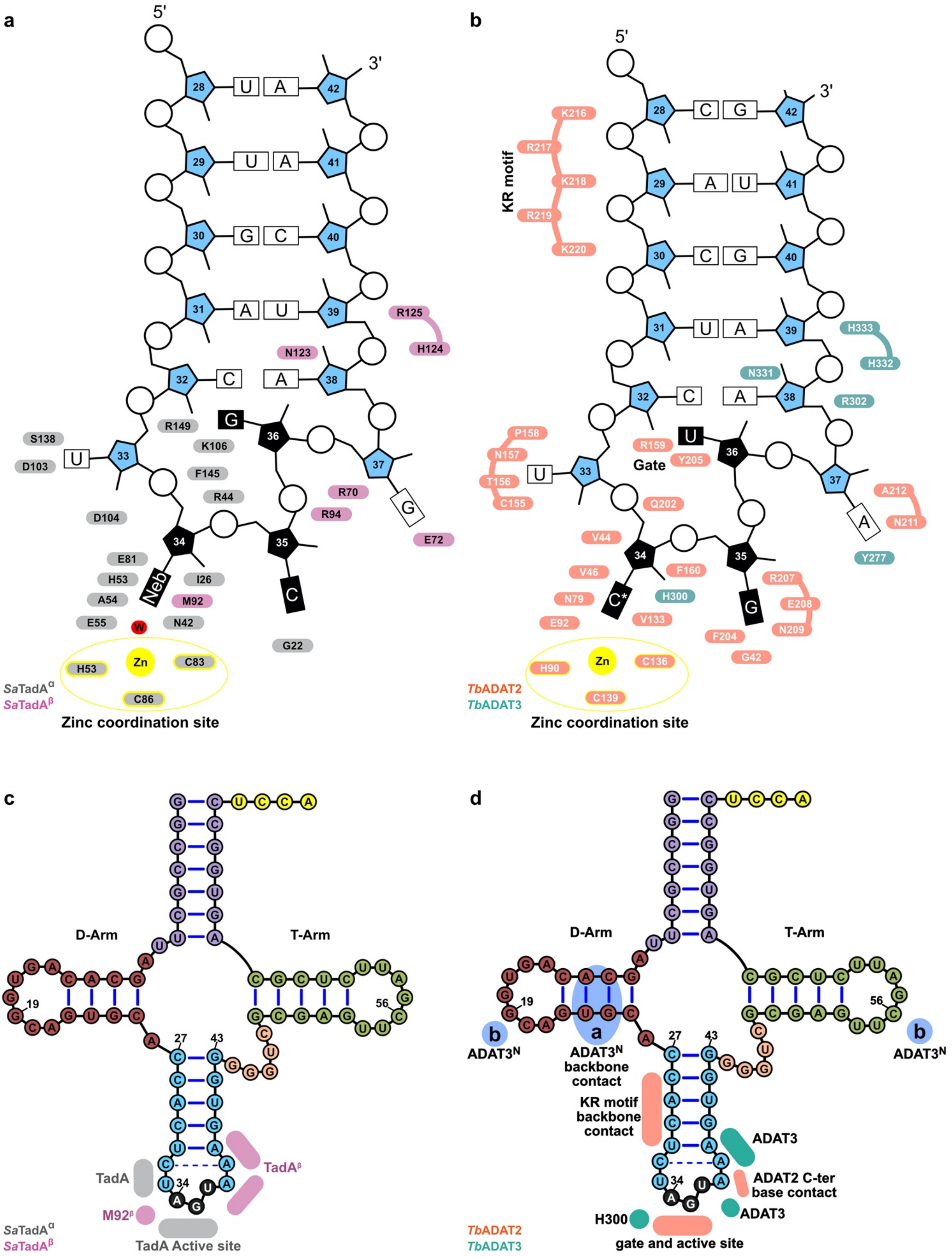

a
b
c
d
5'
3'
Zinc coordination site
Zn
SaTadA^α
SaTadA^β
TbADAT2
TbADAT3
KR motif
Gate
D-Arm
T-Arm
TadA
TadA^β
M92^β
TadA Active site
ADAT3^N
ADAT3^N backbone contact
KR motif backbone contact
ADAT3
ADAT2 C-ter base contact
H300
gate and active site

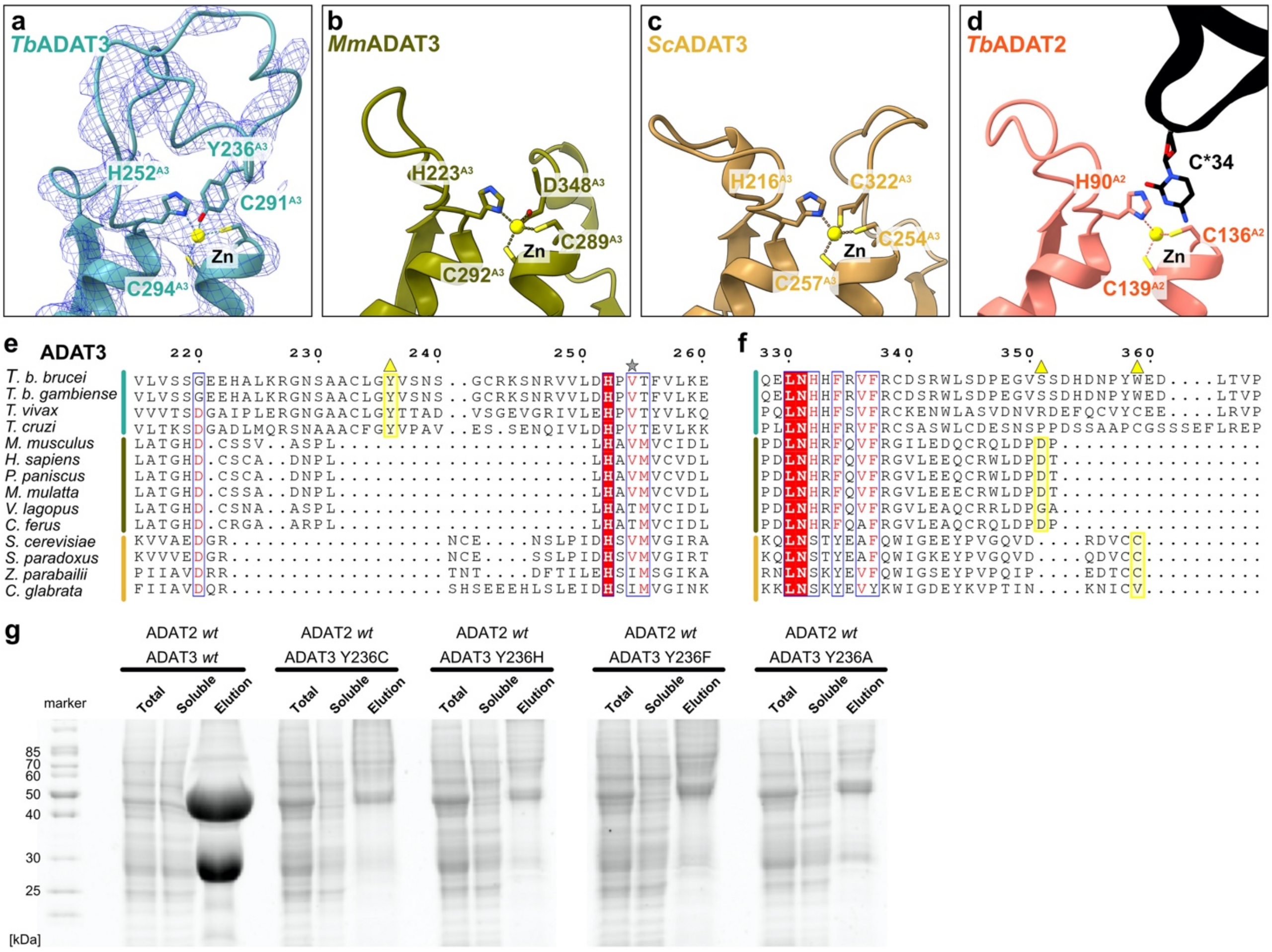
a
TbADAT3
Y236A3
H252A3
C291A3
Zn
C294A3
b
MmADAT3
H223A3
D348A3
C289A3
Zn
C292A3
c
ScADAT3
H216A3
C322A3
C254A3
Zn
C257A3
d
TbADAT2
C*34
H90A2
C136A2
Zn
C139A2
e
ADAT3
T. b. brucei
T. b. gambiense
T. vivax
T. cruzi
M. musculus
H. sapiens
P. paniscus
M. mulatta
V. lagopus
C. ferus
S. cerevisiae
S. paradoxus
Z. parabailii
C. glabrata
f
g
ADAT2 wt
ADAT3 wt
ADAT2 wt
ADAT3 Y236C
ADAT2 wt
ADAT3 Y236H
ADAT2 wt
ADAT3 Y236F
ADAT2 wt
ADAT3 Y236A
marker
Total
Soluble
Elution
85
70
60
50
40
30
25
[kDa]

| **Extended Data Table 1. Summary of Cryo electron microscopy data collection and refinement statistics** | |
|---|---|
| **Data Collection** | tRNA bound *Tb*ADAT2/3 |
| Microscope | Titan Krios |
| Voltage | 300 kV |
| Camera | Quantum-K2 |
| Energy filter | GIF |
| Spot size | 8 |
| Beam size | 650 nm |
| Pixel size (A/pix) | 0.81 |
| Preset defocus range | -1.0 to -1.8 |
| Stage tilt | 30 deg |
| Dose rate ($e^-$/pixel.s) / total exposure ($e^-/A^2$) | 3 / 55 |
| Number of frames per movie | 80 |
| Number of movies | 6,145 |
| **Data Processing** | |
| Initially warp autopick particles | 1,156,506 |
| Final number of particles | 105,718 |
| Resolution FSC | 3.62 |
| Sharpening method | DeepEMhancer |
| EMDB accession number | |
| **Refinement** | |
| PDB accession number | |
| No atoms | 5155 |
| Residues (protein) | 459 |
| Residues (RNA) | 76 |
| $CC_{box}$, $CC_{mask}$, $CC_{volume}$ | 0.69, 0.65, 0.67 |
| R.M.S.D. | |
| Bond lengths | 0.003 |
| Bond angles | 0.651 |
| Ramachandran favored (%) | 94.85 |
| Ramachandran allowed (%) | 5.15 |
| Ramachandran outlier (%) | 0 |
| MolProbity score | 2 |
| Clash score | 13.77 |

**Extended Data Table 2. Summary of published tRNA binding mutants in the ADAT3 N-terminal domain and their corresponding residues in *T.brucei***

| **ADAT3$^{N}$ mutant** | **Effect** | **Corresponding *Tb* residues** |
|---|---|---|
| *Sc*K72A/R73A | tRNA interaction defect (Liu et al., 2020) | K48/R49 |
| *Sc*R75A/K76A | tRNA interaction defect (Liu et al., 2020) | R51/R52 |
| *Mm*K53E/R54E/R61E | Deamination defect (Ramos-Morales et al., 2021). | E24/R25/S32 |
| *Mm*K53E/R54E/R61E/K76E/R82E | Deamination defect (Ramos-Morales et al., 2021). | E24/R25/S32/K48/S54 |
